# Climatic niche evolution of infectious diseases driving amphibian declines

**DOI:** 10.1101/2022.05.13.491758

**Authors:** Gajaba Ellepola, Jayampathi Herath, Sun Dan, Marcio R. Pie, Kris A. Murray, Rohan Pethiyagoda, James Hanken, Madhava Meegaskumbura

## Abstract

Climate change and infectious diseases continue to drive global amphibian population declines, contributing to one of the greatest vertebrate extinctions of the Anthropocene. Currently around 16% amphibian species across the world are affected by four pathogens – *Batrachochytrium dendrobatidis* (*Bd*), *B. salamandrivorans* (*Bsal*), *Ranavirus* and *Perkinsea*. A climatic context behind the dispersal of some of these diseases is hypothesized. However, the interplay between niche conservatism (NC) and climatic niche evolution (CNE), essential to understand disease evolution and dispersal, has so far received little attention. Here we show that the impacts of amphibian pathogens are intensifying as their climatic niches evolve. NC-based analyses suggest that niches of these diseases overlap, especially in Europe and East/southeast Asia (ESEA), and that all four pathogens will continue to devastate amphibians through seasonality shifts and range expansions, penetrating deeper into temperate regions and global amphibian diversity hotspots. *Bd* will spread over diversity-rich mountain ranges and ranaviruses will overwhelm lowlands. CNE-based analyses suggest that the earliest lineages of these diseases originated in colder regions and that some lineages subsequently evolved towards warmer climatic niches. We caution that quiescent, warm-adapted strains are likely to become widespread and novel ranaviruses adapted to local climatic conditions and new hosts are likely to emerge. These results portend the dangers of introducing pathogens into new regions given their ability to adapt to changing climate scenarios. In a climatic background conducive to most of these diseases, frequent monitoring, enhanced biosecurity measures and policy reforms are needed for disease control and mitigation.

## Introduction

Pathogens that cause amphibian diseases, largely unknown only a few decades ago but having found providence through human activities^1,2^, are now dispersing across the world. Working synergistically with other environmental stressors, they are driving amphibian population declines. This sets the stage for possibly the greatest vertebrate species extinction witnessed to date^3-6^. About 1350 of Earth’s 8400 amphibian species are known to have been infected up to now^3,7-11^, with *Batrachochytrium dendrobatidis* (*Bd*), *B. salamandrivorans* (*Bsal*), *Ranavirus* and Severe Perkinsea Infection (SPI) being the most significant (Supplementary Table 1). While these pathogens are associated with population declines, extirpations or extinctions in wild amphibians, chytridiomycosis (both *Bd* and *Bsal*) and ranavirosis are currently recognized as notifiable diseases by the World Organization for Animal Health^12^.

Some of these diseases were first detected by studying the devastation they caused well beyond their native regions of origin. *Bd* was uncovered in Central America and Australia^13^, but its origins are in Asia^5^. By now, numerous lineages have been described, which are known to infect amphibians in 93 countries across the world^10^. *Bsal* was first detected following salamander die-offs in Europe^14^, again with origins in Asia^7^. *Bsal* has not yet been reported in the southern hemisphere, likely due to the near absence of its host taxon (salamanders). *Frog virus 3*, the first ranavirus isolate, was discovered in the United States^15^ but has now been recorded across 25 countries covering all continents except Antractica^8^. Several clades of cryptic SPI have been detected in five countries in Africa, Europe and North America^3,16^ since its discovery in the United States^17^. It is an emerging frog disease in North America^18^ of unknown global concern. Despite their widespread impacts, the patterns of distribution and dispersal of these diseases remain poorly understood, limiting the formulation and implementation of adequate disease control measures in affected or at-risk regions.

These diseases have dispersed beyond their northern-hemisphere centers of origin^5,7,8^ and for the most part their spread has been human-aided and potentially facilitated by recent climate change. Several studies highlight the role of climate in providing the background for the distribution and impacts of amphibian pathogens^19-24^. Yet, and despite the urgent need for mitigation and the clear potential role of climate change to facilitate worldwide expansion, global projections of future spread of amphibian diseases remain poorly resolved.

The contrasting theories of niche conservatism (NC)^25^ and climatic niche evolution (CNE)^26,27^, or the interplay between NC and CNE^28^, have been central to efforts to understand the distribution and dispersal patterns of species. However, few studies have applied such theory to invasive pathogens that contribute to catastrophic biodiversity loss^29,30^. NC proposes that species maintain a relatively stable ecological niche over time, particularly with respect to climatic axes. In contrast, CNE postulates that some lineages adapt to novel niches while others remain in climatic stasis, i.e., close to their ancestral climatic niche.

To better understand the distribution and dispersal patterns of these four pathogens, we carried out a series of analyses based on NC and CNE in a phylogenetic context. We examined the climatic niches of these pathogens as a proxy of their n-dimensional environmental niches. We also contrast NC and CNE concepts to examine potential evolutionary shifts in climate preferences that may help explain current—and predict future—geographic spread. We use global occurrence data and bioclimatic variables, in combination with spatial point process models^31^ and maximum entropy (Maxent) models^32^, to assess the climatic suitability for these pathogens and the extent to which they currently occupy their available climatic niche spaces and their corresponding potential global geographic distributions. We then quantify their climatic niche preferences based on a principal components analysis. Further, we use CMIP6 downscaled climatic scenarios (2021-2040 time period) to predict future potential geographical range shifts due to a changing climate and evaluate the extent to which biodiversity hotspots may eventually be infiltrated by these diseases.

The above analyses begin by assuming NC, wherein species are thought to maintain their climatic niche preferences over time. Thus, in the future, species are predicted to be distributed in areas that have climatic conditions similar to those they currently occupy^25^. In addition, and to evaluate the possibility of CNE, we hypothesize that some variants/isolates may have deviated from their ancestral climatic niches to achieve novel climate characteristics. We assume that most variants/isolates remain in climate stasis close to the climatic conditions in which they evolved, such that the deviation of some variants/isolates from the inferred ancestral niche (taken as the average of all strains/lineages^26,33^ serves as an indicator of climatic niche evolution. Such deviation is visualized by means of traitgrams, disparity-through-time plots and bin-based ancestral reconstructions. In total, this comparative analysis reveals the climatic context that underpins the distributions of these four amphibian diseases. It also allows both comparisons among pathogens and an examination of the geographic shifts that could be expected under future climatic projections. This, in turn, enhances the critical knowledge base relevant to disease mitigation.

## Results and Discussion

### Predicted distributions of amphibian diseases

We found that the climatic niches of these four pathogens, and hence their predicted geographic distributions, overlap considerably, especially in Europe and East/Southeast Asia (ESEA) (Fig. 1). *Bd* has a potential geographic distribution that includes North America, South America, regions bordering the Mediterranean, southern Africa, ESEA and eastern Australia. Meanwhile, the potential distribution of *Bsal* includes Europe, the Mediterranean, and ESEA. Importantly, whereas the predicted distributions of *Bd* and *Bsal* are consistent with previous reports^10,29,30^, ours is the first study to predict in addition the global distribution of ranaviruses and SPI. Ranaviruses have a predicted potential distribution that spans Central America, Europe, ESEA and Australia, while SPI is predicted to be potentially distributed around eastern North America and parts of South America, Europe, Africa, ESEA and Australia.

**Fig. 1.**
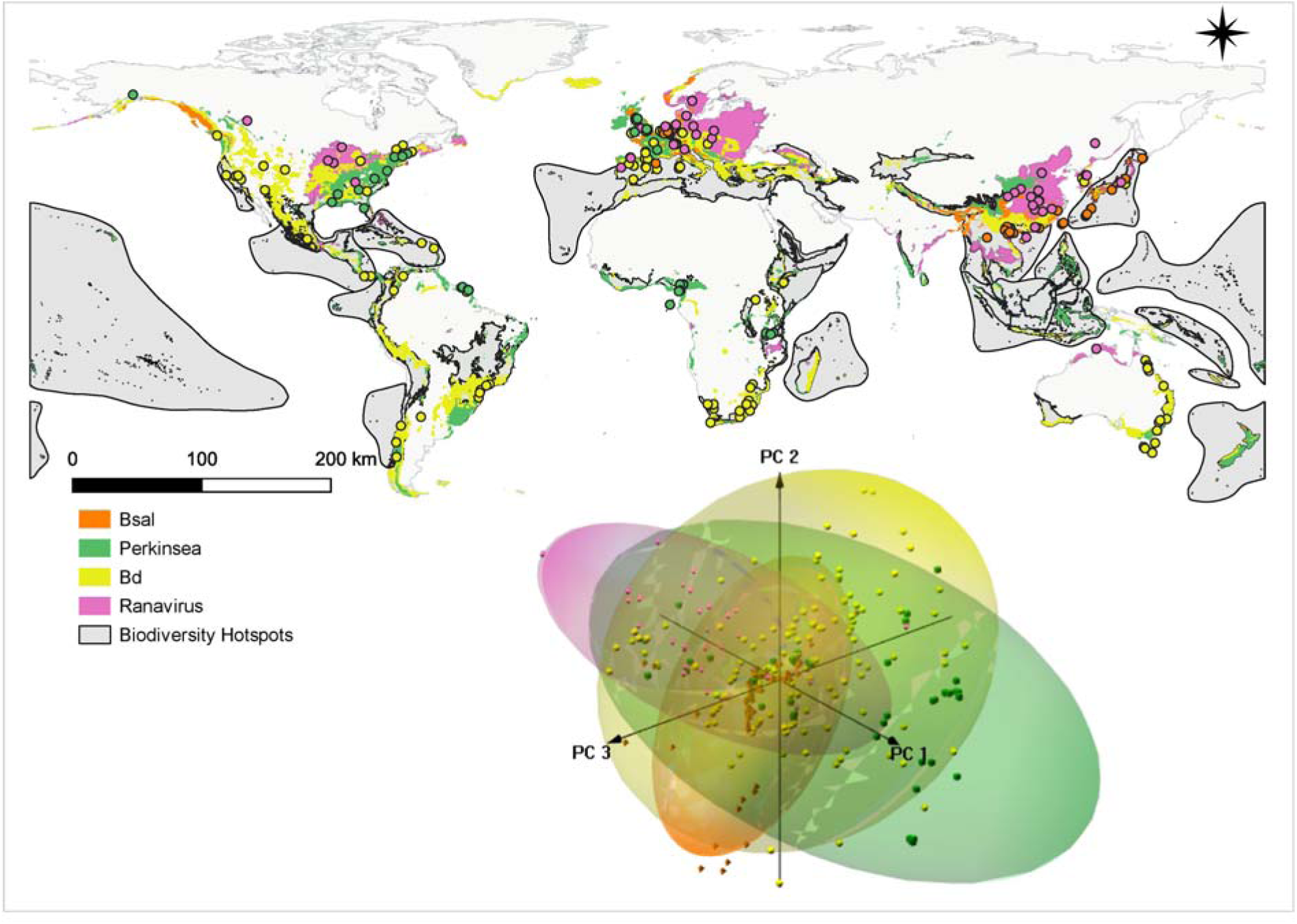
Potential geographic distributions of *Bd, Bsal*, ranaviruses and *Perkinsea*, and their climatic niche spaces defined by the first three principal components. Disease distribution range polygons are based on > 0.5 Maxent probability distributions (see Extended Data Figure 1for broadly congruent PPM results). Colored dots indicate up-to-date occurrence records of these four amphibian diseases. Their potential distributions largely overlap, especially in Europe and ESEA. *Bsal* (possibly due to a dearth of salamanders, a principal host taxon) has not been reported in the southern hemisphere. Most occurrences are currently seen outside biodiversity hotspots (shaded in grey). The 3D PCA plot indicates climatic niche space defined by the first three principal component axes of climatic niches of these diseases. Each point represents the average climatic conditions for each taxon/variant/strain. PCA loadings are provided in Supplementary Table 2. The first three axes explain more than 75% of the variance in the data. Temperature dependence of each disease is indicated by the PCA loadings, in which PC1 (45.7%) is represented mainly by ‘Mean annual temperature of coldest quarter, Annual mean temperature, Min temperature of coldest month’; PC2 (20.5%) by ‘mean diurnal range’; and PC3 by ‘Temperature seasonality and Precipitation of warmest quarter’. Climatic niches of the four diseases overlap to a certain extent but segregate along PC2 and PC3 (Supplementary Table 3). *Perkinsea* occupies the broadest climatic niche space, while *Bsal* occupies the narrowest.

The analyses also identify climatically suitable areas that are not yet infiltrated by these diseases. These include many amphibian-rich Global Biodiversity Hotspots^34^ (Fig. 1, map). In general, both spatial point-process models (PPM; Extended Data Figure 1) and Maxent distribution models (Fig. 1) yield similar results. However, PPMs generally yield lower probability-of-occurrence values, possibly due to edge effect and spatial dependence^31^.

In a PCA of the average bioclimatic conditions associated with occurrence of the studied taxa, the first three axes explain over 75% of the data variance (Fig. 1 sub panel, Supplementary Table 2). For PC1 (45.7%; mainly bio11, bio1, bio6; Supplementary Table 2), higher scores correspond to less-cold-tolerant variants/isolates and *vice versa*. For PC2 (20.5%; mainly bio2), higher scores represent more-temperate variants/isolates while lower scores represent more-tropical ones. Consistent with previous studies, these results confirm that all four pathogens are temperature-sensitive^14,18,35,36^. More specifically, they indicate that the pathogens’ distributions are primarily constrained by temperature, with other climatic factors being of secondary importance. Similar to the geographic overlap among pathogens based on niche models and PPMs (Fig 1, map), the climatic niche spaces of the four pathogens also overlap to a considerable degree (Fig. 1 sub panel, Fig. 2 main panel). In general, higher probabilities of occurrences of all four diseases occur towards cooler, subtropical climatic conditions, especially in the case of *Bsal* and ranaviruses. *Bd* and SPI extend more widely into the tropics, i.e., regions of higher temperature (Fig. 2 main plot) ^3,37,38^. Distribution models for both *Bd* and SPI further suggest that these pathogens could potentially be distributed in cool, high-altitude areas in tropical regions as well (Extended Data Figure 2).

**Fig. 2.**
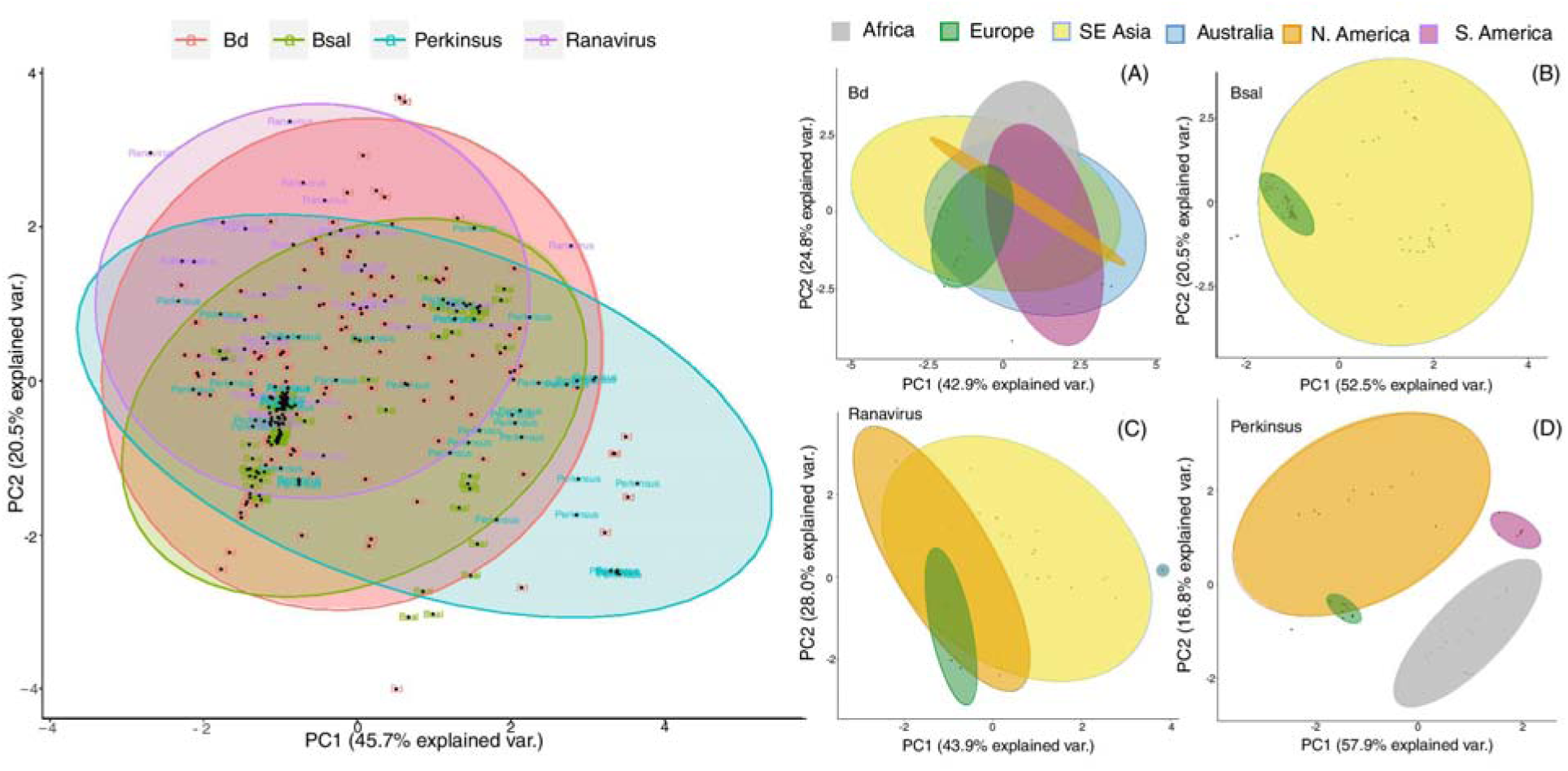
2D climatic niche spaces traced by the geographic distribution regions of *Bd, Bsal*, ranaviruses and Perkinsea. Ellipses outline 95% confidence intervals. There is a higher prevalence of all four diseases under colder climatic conditions. Generally, *Bsal* and ranaviruses seem to prevail under cold and subtropical climatic conditions, whereas *Bd* and Perkinsea are more widely distributed from warm to cold climatic conditions. *Bsal* is represented by two distinct climatic clusters that segregate along PC1 (main plot). This reflects the widely disjunct distribution of the introduced European population vis-à-vis the native Asian population. Geographic regions of each disease mapped on climatic space (Figs. 2A–D) identify these two clusters as *Bsal* in Europe (colder climates) and *Bsal* in ESEA (tropical/subtropical climates). The climatic niche space of *Bsal* in Europe is more conservative than that of *Bsal* in ESEA and is found at the edge of the latter’s niche space. *Bd* in North and South America has a restricted climatic niche along PC1 but a broader occupation in ESEA and Australia. Ranaviruses in ESEA are represented by broader climatic niches, whereas Perkinsea in different regions occupy non-overlapping climatic niches, possibly due to inadequate detection of this emerging pathogen.

### Breadth of the climatic niches of amphibian pathogens

SPI and *Bsal* have the broadest and narrowest climatic niche space occupations, respectively (Fig 2, main plot). Yet, the relatively narrow climatic niche of *Bsal* appears to consist of two different climatic clusters, one that is a restricted subset within the 95% confidence limits of the other (Fig. 2, main plot, Fig. 2B). When stratified by geographic region, these clusters are associated with distinct clusters of *Bsal* in Europe (colder niche) and *Bsal* in ESEA (relatively milder climates), which represent widely disjunct distributions in the introduced European population and the native Asian population^7^. The *Bsal* introduced to Europe occupies a considerably restricted climatic niche space than does native *Bsal* in ESEA, possibly due to incomplete invasion in Europe. *Bd* in Africa, South America and Europe has a restricted climatic niche along PC1 but a broader occupation in North America, ESEA and Australia (Fig 2A). Ranaviruses in ESEA are represented by broader climatic niches, whereas SPI in different regions occupies non-overlapping climatic niches, an artifact possibly due to sparse sampling of this emerging disease^3,18,39^. Overall, *Bd, Bsal* and ranaviruses tend to occupy a broader climatic niche in Asia relative to other regions, which suggests that strains/variants in Asia are generalists that have been able to adapt to varying climatic conditions and/or that they occupy a greater potential niche space as some originated in Asia (*Bd* and *Bsal*). In this respect, variants in Asia may be of particular concern in terms of risk of establishment when introduced to novel regions (e.g., through live-animal trade) due to their higher adaptability to different climatic conditions.

Climatic niche axes differ in some respects among the four pathogens (Extended Data Figure 3). Although temperature variables limiting cold tolerance are common to all the pathogens along the PC1 axis, other variables differ along the PC2 and PC3 axes (Supplementary Table 3). Temperature sensitivity (explained by PC1 for all pathogens) is confirmed generally by the fact that *Bd* and *Bsal* are reported during milder weather or during cooler months in warm regions^36^, whereas ranavirosis and SPI in the United States is reported in warmer months^18,35^. *Bsal* and SPI seem to be dependent on rainfall during the driest period, whereas *Bd* and ranaviruses seem to be less dependent on rainfall on PC2 but require less variable diurnal temperatures (isothermal conditions) along PC3 (Extended Data Figure 3). Overall, seasonality and precipitation variables appear to segregate the climatic niche spaces of these diseases. The dependence of *Bsal* and SPI on rainfall during the driest periods can be inferred from the life histories of their potential hosts. *Bsal* affects salamanders and newts^14,40^ and SPI affects post-metamorphic stages of anurans^18,39^. The life histories of these hosts are largely dependent on water bodies, with associated differences in disease spread^39,41,42^. Transmission remains low during dry periods but may increase with the onset of rains, which usually coincides with the hosts’ breeding periods as well.

**Fig. 3.**
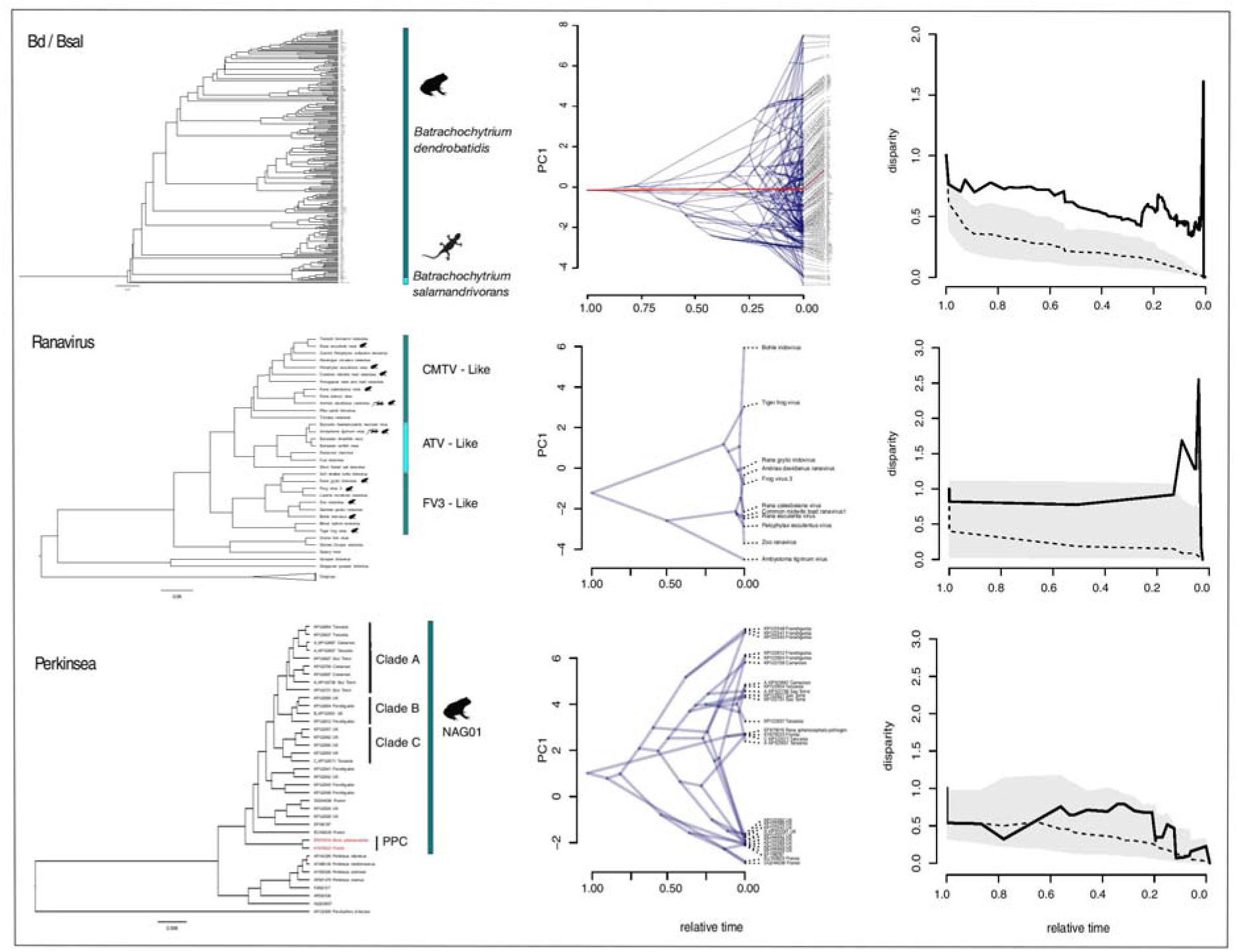
Molecular phylogenies (left), variation in the rate of evolution of the climatic niche (middle) and relative disparity-through-time plots (right) of pathogens causing major amphibian diseases (*Bd*/*Bsal*, ranaviruses and Perkinsea) worldwide. Phylogenies were reconstructed using available genomic and Sanger sequence data and represent the most current phylogenies of the variants/strains of the four main amphibian disease-causing agents (left column). *Bd* and *Bsal* phylogenies include 235 *Bd* variants and a single *Bsal* variant, whereas the ranavirus phylogeny includes 33 strains representing CMTV-like, ATV-like and FV3-like ranaviruses. Eleven ranaviral strains known to infect amphibians are indicated by silhouettes of frogs and salamanders. The Perkinsea phylogeny is represented by 29 NAG01 variants. The variants highlighted in red indicate two representatives from the pathogenic Perkinsea clade (PPC). Deviations from the ancestral climatic conditions of the major amphibian disease-causing pathogens are explained by the first principal component axis (middle column). The X-axis depicts relative age from the origin of the pathogen to the present; nodes indicate the inferred climatic niches for the most recent common ancestor of the extant taxa defined by that node. Among all pathogens, some variants/strains have evolved from their ancestral climatic conditions towards a colder climate, while most others have evolved towards a warmer climate. *Bsal* has not deviated much from its original climatic condition, despite occupying new habitats in Europe. A cold-adapted ancestral climatic state is recovered for ranaviruses; divergent branches are more frequent near the present, suggesting recent climatic niche evolution. Perkinsea distributions suggest a pattern of climatic niche evolution towards two temperature extremes along PC1, but this may be an artifact of inadequate sampling. Relative disparity-through-time (DTT) plots of PC1 scores depict change of rate of climatic niche evolution of the four major amphibian disease-causing pathogens (right column). Solid lines indicate the observed DTT, whereas dashed lines and corresponding polygons represent the averages and 95% confidence intervals of the expectations given a constant accumulation of disparity over time based on 999 pseudo-replicates. For *Bd*, closely related variants differ considerably in their climatic niches. A higher rate of climatic niche evolution is observed in *Bd* throughout its evolutionary history; for ranaviruses, the acceleration along PC1 seems rapid and recent. The disparity of climatic niche evolution for SPI did not deviate much from a null model of constant accumulation of disparity over time.

### Climatic niche evolution (CNE) of amphibian pathogens and their clades/strains

Our climate niche-based analyses provide evidence for potential CNE in various strains/isolates. The CNE traitgrams of the four pathogens based on ancestral reconstructions of PC scores of the climatic niche axes (the evolving trait) suggest that all European and a few North American variants of *Bd* have evolved from their ancestral climatic conditions towards a colder climate. Meanwhile, most other *Bd* variants suggest possible evolution towards a warmer climate (Fig. 3). Here, “ancestral climatic conditions” refers to the PC value at the node of the phylogenetic tree. The alternative bin-based method, which explicitly incorporates phylogenetic uncertainty in ancestral reconstructions, also supports possible CNE in different *Bd* strains (Extended Data Figure 4). For example, strain ASIA-1 shows potential niche expansion under bio2, bio3, bio5, bio11, bio17 and bio19, while niche retraction is evident in CAPE under most of the climatic variables. Moreover, according to the DTT plots, the climatic niche of *Bd* evolves at a faster rate than a null model of constant accumulation of disparity over time. In contrast, based on its traitgram, *Bsal* does not appear to have deviated substantially from its ancestral climatic conditions (subtropical and temperate climates in Asia) despite occupying novel habitats in Europe (Fig. 2). The bin-based ancestral reconstruction method, however, shows potential evidence for niche retraction under bio3, bio6 and bio11 (Extended Data Figure 4). These results indicate that the climatic niche of *Bsal* is conservative. Hence, the broader climatic conditions under which it currently occurs likely reflect the conditions under which it evolved^7,43^.

**Fig. 4.**
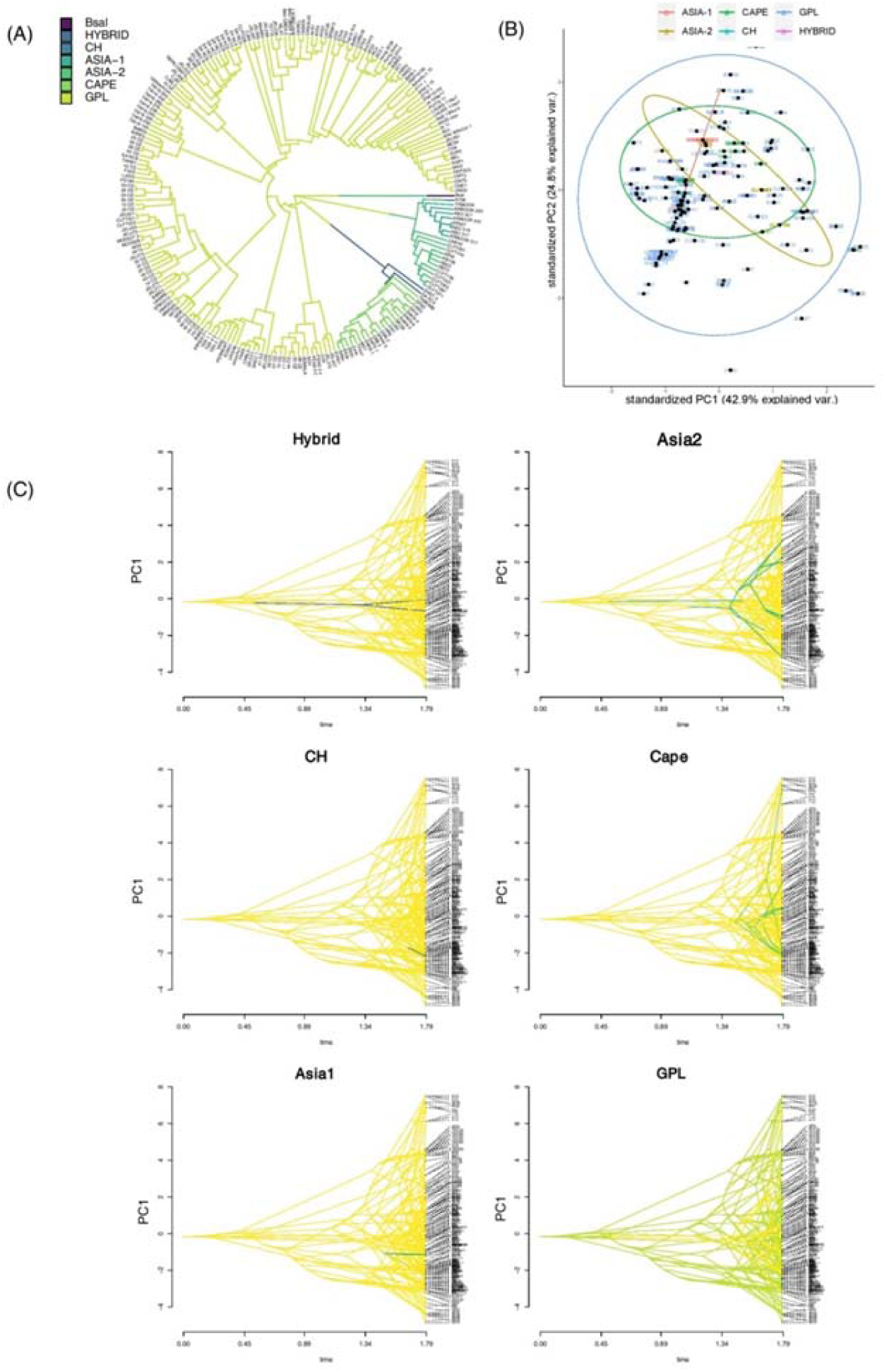
Climate correlates of different *Bd* lineages. (A) Six main lineages of *Bd* (GPL, CAPE, ASIA-1, ASIA-2, CH and HYBRID) and *Bsal* traced on the phylogeny. *Bsal*, ASIA-1, ASIA-2 CAPE are early diverging lineages and GPL is the most recently diverging lineage. (B) Climatic niche occupation of the six main lineages of *Bd*. Circles represent 95% confidence intervals. GPL exhibits the broadest niche occupation, followed by CAPE and Asia2. All other lineages are found within the broad climatic niche space of GPL. Comparison among the lineages in (A) and (B) suggests that GPL, the most recently evolved lineage, has evolved the broadest climatic niche space. (C) The six main lineages of *Bd* traced on the traitgram showing climatic niche evolution of *Bd* lineages. Yellow branches represent the background data whereas the green branches represent the lineage being considered. GPL shows the greatest deviation from the ancestral climatic niche. CAPE and Asia2 also show considerable deviation from the ancestral climatic niche, whereas ASIA-1, CH and HYBRID have retained it.

For ranaviruses, the ancestral climatic conditions inferred from traitgrams predict cold adaptation, which is different from most current ranavirus strains. A few strains have evolved towards warmer climates, and some have evolved towards cooler conditions, but only recently in their evolutionary history (see relative time axis on Fig. 3 center panel). The bin-based ancestral reconstructions for different clades of ranaviruses supply potential evidence for niche retraction in ATV-like ranaviruses under bio11 and bio15, and for CMTV-like ranaviruses under bio3. Potential niche expansion is evident for CMTV-like ranaviruses under bio6, bio14 and bio17, while no evidence for CNE is apparent in FV-3-like ranaviruses (Extended Data Figure 5). However, in a CNE context, pathogens would require time to establish (ranaviruses may be an exception). Climatic niche evolution of SPI seems to be slow and irregular. However, this pathogen seems to have evolved towards two temperature extremes along PC1, with cold-adapted variants concentrated in areas with high precipitation along PC2. While the absence of intermediate variants may be an artifact of sparse sampling, the potential niche expansion shown by the Pathogenic Perkinsea Clade along bio2, bio6, bio14 and bio17 could be of particular interest (PPC; Figs. 3, Extended Data Figure 6). Nevertheless, because some variants/strains among these pathogens are adapted to warmer climatic conditions, we infer that these pathogens undergo CNE. This portends that some variants will evolve in response to ongoing climate change. If they do, the vulnerability of cool□ and warm □adapted hosts could escalate as the pathogens evolve their climatic niches towards unusually warm and cool temperatures, respectively^44,45^.

**Fig. 5.**
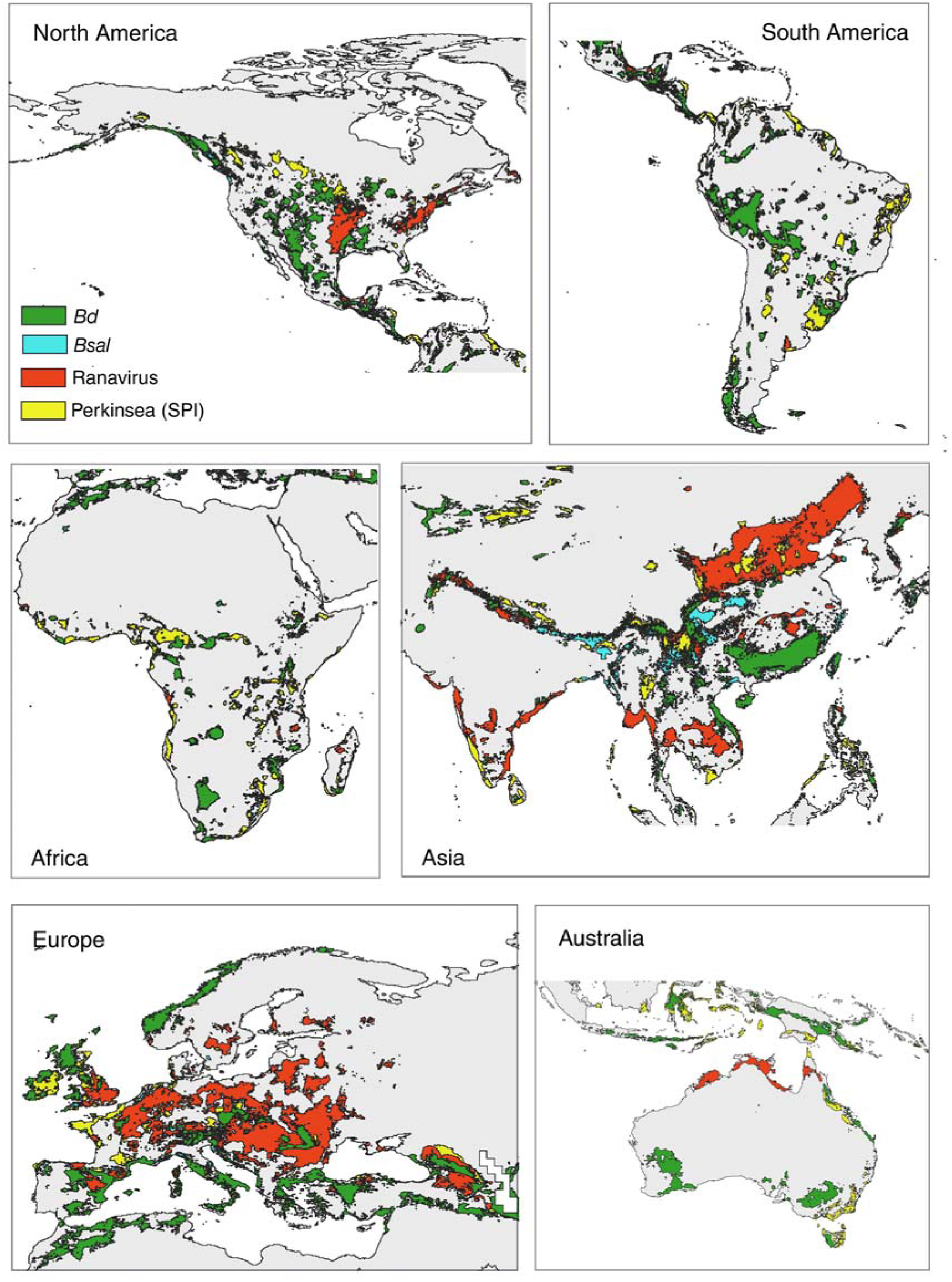
Predicted potential mean range expansion (in 2040) of *Bd, Bsal*, ranavirus and Perkinsea based on CMIP6 downscaled future climate projections. An expansion of climatic conditions suitable for *Bd* is evident throughout the subtropics and around the Mediterranean climatic region, while *Bsal* continues to expand specifically in high-altitude regions in Asia. Ranaviruses are evident throughout the tropics and subtropics. Perkinsea will expand into Asia, to Africa, Europe, South America, the isthmus of Panama, North America and some parts of Indonesia in another 20-30 years. See Extended Data Figure 8 for areas that are currently suitable but likely to become less suitable under future climatic scenarios.

### Climatic context of Bd virulence

With available data for *Bd*, the best known of the four pathogens, we further investigated whether climatic niche differences exist between lineages that differ in their apparent virulence. Six known virulent *Bd* lineages (GPL, CAPE, ASIA-1, ASIA-2, CH and HYBRID)^5^ were traced on our PC plots and traitgrams (Fig. 4). GPL, the most recently evolved and pandemic-causing lineage that is primarily responsible for global disease-associated amphibian declines and extinctions, exhibits the broadest niche occupation, followed by CAPE (also frequently disease causing) and ASIA-2 (rarely observed to cause disease in wild amphibians). All other lineages fall within the broad climatic niche space of GPL (Fig. 4B). Traitgrams also suggest that GPL has deviated the most from its ancestral climatic niche, followed by CAPE and ASIA-2. However, ASIA-1, CH and HYBRID seem to have retained the ancestral climatic niche (Fig. 4C). These results do not offer clear evidence that virulence has a strong climatic niche component at the lineage level (Extended Data Figure 7).

### Future climatic change and amphibian pathogen distributions

Predicted geographic distributions of the four major amphibian diseases two decades from now (2040) suggest that *Bd* and *Bsal* will occupy less of their overall current potential distributions while ranavirus and SPI will likely occupy more of theirs (Figs. 5, Extended Data Figure 8). For *Bd*, this prediction is corroborated by earlier findings^19^. However, if appropriate host species are present, *Bd* still has the potential to expand into high-elevation areas in North America, the Andes Mountains in South America, parts of South Africa, Asia, Europe, southern parts of Australia and the Sunda Islands in response to a changing climate (Extended Data Figure 2). *Bsal* is predicted to spread more widely into high-altitude areas in temperate regions but remain largely absent from tropical regions. The overall reduction of *Bsal* seen in the projections may be explained by its apparently narrow and conservative, cool-adapted climatic requirements in the context of global warming. In contrast, ranaviruses appear likely to expand their distribution throughout the tropics and subtropics in regions such as North America, Europe, South Asia, ESEA and northern parts of Australia. Similarly, apart from its current distribution, SPI may also expand into Asia, Africa, the Panamanian isthmus, and some parts of Indonesia if given opportunity to establish. Despite the seemingly complacent scenario regarding future expansion of the global strain of *Bd* and ranaviruses, the situation remains precarious, especially in relation to biodiversity hotspots and tropical montane regions. Furthermore, potential hosts that are adapted to relatively cool conditions may be the most vulnerable to increases in mean temperature^44^. We therefore predict that Asian amphibians will soon be at greater risk due to these diseases. Among the four pathogens, ranavirus, with its potential to exploit even non-amphibian host species, will become widespread in temperate and tropical regions, while infiltrating biodiversity hotspots. These pathogens thus pose an emerging, perhaps imminent threat to amphibians throughout the world^8^.

### Potential global risks of amphibian diseases

In the absence of adequate mitigating measures, these diseases are likely to further devastate amphibian populations. The wide availability of additional regions that are suitable for the establishment of these diseases and their presence in the commercial trade of amphibians^46^ calls for regulations that prevent, or at least inhibit, further anthropogenic spread. Biodiversity hotspots and high-elevation areas with high amphibian diversity are especially vulnerable^47,48^. There is mounting evidence that species popular for frog aquaculture, such as the American bullfrog (*Lithobates catesbeianus*) and the Pig frog *(L. grylio*) imported from North America to Asia, may have acted as reservoir hosts for ranavirus strains^49^. SPI outbreaks are causing mass mortalities of tadpoles across the United States since the pathogen was first detected in 1999^18^ and are likely distributed throughout much of the Nearctic region^18^. Perkinsea SSU rDNA sequences obtained from seemingly healthy wild tadpoles sampled in Panama and from captive-bred *Hyla arborea* tadpoles from the United Kingdom are similar to Pathogenic Perkinsea Clade sequences sampled from mortality events in the United States^16^. However, *Bd* and *Bsal* likely originated in Asia^5,7^. We predict that amphibians in many parts of the southern hemisphere are vulnerable to all four diseases as they have never encountered the corresponding pathogens until recently, if at all. Furthermore, introduction of pathogens from Asia to other regions may pose serious threats to amphibians worldwide because they have the ability to persist under broader climatic tolerance limits, especially if climatic niches evolve with time. Hybridization among variants of a particular pathogen species can result in the emergence of lineages adapted to warmer conditions as well as enhanced virulence. Variants already adapted towards warmer climates require especially close monitoring.

While several mitigatory strategies have been developed (mostly in vitro; Supplementary Table 4), these are difficult to implement in the wild, which leaves prevention of dispersal as the most potent form of mitigation available at present. At a global scale, climate change alone is likely insufficient to instigate the spread of these amphibian pathogens into vulnerable but currently unoccupied areas; human-aided spread remains critical. This provides both challenges and opportunities to controlling disease spread, as biosecurity measures are easier to implement than is ending climate change. Nevertheless, controlling disease spread remains a profound challenge, where distances are not barriers for human-induced transmission, in the absence of strong and enforceable biosecurity measures. Our results draw attention to the likely serious consequences of introducing pathogens into novel regions, while the exacerbating effect of climate change in promoting pathogen establishment and spread appear to outweigh any gains in terms of controlling disease impacts. In situations where amphibians are transferred across national borders for commercial purposes, International Import Risk Analysis (IRAs) should be used to establish or revise trade or translocation guidelines for wildlife^50^. Risk analysis on wildlife species in trade, pre-border pathogen screening, and voluntary support could reduce the substantial costs associated with invasive species as well as protect public, environmental and animal health^52^. Unless adequate mitigation measures are taken, these diseases could ultimately devastate already imperiled amphibian populations. Effective biosecurity measures, better monitoring and policy innovations are crucial for reducing the otherwise inevitable destruction that will result from their global spread.

## Methods

### Phylogenies

Initially, we collected all available data on different aspects of the four amphibian diseases of interest and prepared a summary table of their prominent characteristics (Supplementary Table 1). This allowed us to identify all potential variants/strains and isolates of these diseases described in literature. Extensive taxon sampling is important in phylogenetic systematics, as it increases accuracy and support of evolutionary relationships^51^. Hence, we present updated phylogenies of the taxa being considered. The available genetic data for different lineages is uneven, from species with none to ones in which Sanger sequence data and whole genomes have been reported. To derive phylogenies with complete taxon sampling that combine all available data, we used a two-staged Bayesian approach named PASTIS^52^, which uses as inputs a backbone topology based on molecular data, a set of taxonomic postulates (e.g., constraining species to belong to their closest sister taxa, specific genera or families), and user-defined priors on branch lengths and topologies. Based on these components, PASTIS produces input files for MrBayes 3.2.5^53^, which generates a posterior distribution of complete ultrametric trees that capture uncertainty under a homogeneous birth-death prior model of diversification and placement constraints. We used PASTIS version 0.1-2, with functions from the APE 5.4^54^ and CAPER 0.2^55^ packages.

To infer the phylogenetic relationships among *Bd* variants, we used the previously published phylogeny of^56^ available at https://microreact.org/project/GlobalBd (last accessed on March 2021). Microreact provides an updated robust phylogeny constructed from genomic data. To infer the phylogenetic relationship between *Bd* and *Bsal*, we used the PASTIS approach and used the *Bd* phylogeny of^56^ as the backbone tree. Next, we used the available genomic data for both *Bd* and *Bsal* from GenBank (last accessed on March 2021) and compiled a data set comprised of 4660 base pairs (bp), which was aligned using MAFFT^57^. The data were constrained based on their sister taxa relationship and was run on MrBayes 3.2.5^53^ under a homogeneous birth-death prior model of diversification for 10 million generations. Convergence was assessed by inspecting the log-output file in TRACER v.1.6^58^ and by ensuring ESS values were above 200. The first 10% of the trees were discarded, and the post burn-in trees were used to infer the maximum clade credibility tree using TREEANNOTATOR v.1.10.4^59^.

A similar method was adopted to infer the phylogenetic relationships among different ranaviral strains. Initially, we retrieved 25 complete genomes of different ranaviral strains from GenBank (last accessed on March 2021), which consisted of more than 100,000 bp. Downloaded nucleotide sequences were aligned using MAFFT^57^ in the Cipres Science Gateway Server^60^. The sequence alignment was analyzed in several tree building programs such as IQtree^61^, NDtree^62^ and CSI phylogeny^63^ by assigning TVMe+R2, the best-fitting nucleotide model. The same topology was recovered regardless of the tree building method; we used the ultrametric tree output from MAFFT as the backbone tree for subsequent analyses. To incorporate additional taxa having only short sequence data into the backbone tree, we downloaded from Genbank nucleotide sequence data for several loci available for all ranaviral taxa (MCP, DPG, RDRASPG and RDRBS). The nucleotide sequences were aligned in MEGA X^64^ and the concatenated file was analyzed in BEAST v.1.4^58^. This step was carried out especially to infer the sister-taxon relationships among ranaviral strains and assign phylogenetic constraints to them, as we could not recover these relationships from published literature. We could retrieve possible sister taxa relationships of the strains that had only partial sequence data. Using these recovered relationships as a guide, we constrained them as ‘soft constraints’ into specific clades and ran the analysis in MrBayes 3.2.5^53^ for 60 million generations under the PASTIS tree building framework^52^. We pruned the resulting maximum clade-credibility tree by removing taxa that do not cause diseases on amphibians and used the resulting tree in subsequent analyses.

To infer phylogenetic relationships among Perkinsea, we downloaded NAG01 sequences deposited by^3^ and^18^in GenBank (last accessed March 2021). Reference sequences for four *Perkinsus* species (*P. atlanticus, P. mediterraneus, P. andrewsi* and *P. merinus*) and a reference sequence for *Parvilucifera infectans* were also used to root the tree. After carefully curating the sequences, an alignment of the sequence data with 641 bp was generated using MUSCLE in MEGA X^64^. Maximum likelihood and Bayesian methods were performed using RAxML-HPC236 and BEAST v.1.4, respectively, through the CIPRES Science Gateway^60^. For the maximum likelihood analysis, a general time-reversible model with gamma distribution was used and 1000 bootstrap iterations were performed. For the Bayesian analysis, a GTR+I model with gamma distribution was used, and the number of generations was set to 5 million. Convergence was assessed by inspecting the log-output file in TRACER v.1.6^58^ and by ensuring ESS values were above 200. The first 10% of the trees were discarded and the post burn-in trees were used to infer the maximum clade credibility tree using TREEANNOTATOR v.1.10.4^59^. However, we used the ultrametric trees obtained from BEAST for further analyses.

The inferred phylogenies (Fig. 3 left column) were congruent with the latest available phylogenies of these four diseases^3,5,18,56,65^. Our *Bd/Bsal* phylogeny includes 235 *Bd* variants and the single known strain of *Bsal*. The *Bd* variants have a wide geographic distribution, which ranges from North America to South America, Europe, Africa, ESEA and Australia, thus infecting frogs and toads worldwide. *Bsal* is distributed in Europe and ESEA and thus infects salamanders in these regions. Seven species of *Ranavirus* have been described according to the updated classification of the International Committee on Taxonomy of Viruses (ICTV): *Ambystoma tigrinum* virus (ATV), *Common midwife toad virus* (CMTV), *Epizootic haematopoietic necrosis virus* (EHNV), *European North Atlantic ranavirus* (LfRV), *Frog virus 3* (FV3), *Santee-Cooper ranavirus* and *Singapore grouper iridovirus* (SGIV)^66^. Ranavirus isolates are considered members of the same viral species if they share > 95% amino acid identity. However, *Ranavirus maximus, Cod iridovirus* and *Short-finned eel virus* are potential new species that remain unclassified^66^. Among these, 11 strains are known to infect amphibians distributed in Canada, China, Japan, Australia, Netherlands, England, Spain, Germany, Italy, Sweden, Slovakia, Poland and the Czech Republic. FV3, ATV, and BIV are recognized species of ranavirus that are known to infect amphibians, though several additional ranaviral strains have been isolated from amphibians^8^ FV3s are amphibian specialists, whereas ATVs are predominantly fish specialists that switched once to caudate amphibians^67^. The Perkinsea phylogeny is represented by 29 NAG01 variants spanning Africa, French Guinea, the United Kingdom, France and the Pathogenic Perkinsea clade (PPC) in the USA.

### Climatic and distribution data

We obtained from published literature present-day geographical occurrences in the form of GPS coordinates for all taxa represented in our three phylogenies of amphibian diseases (*Bd*/*Bsal, Ranavirus* and Perkinsea). *Bd* occurrence data were obtained directly from https://microreact.org/project/GlobalBd^56^. *Bsal* occurrence records were obtained from^7,30,40,43,68,69^. Ranavirus occurrence data were obtained from relevant literature. Occurrence data for Perkinsea were obtained from^3,18^. However, when GPS coordinates were not available for certain taxa (especially ranaviruses and Perkinsea), we arbitrarily assigned a GPS location based on the name of place (village/town/county/province) provided in published literature, assuming that average climatic conditions are similar among nearby locations. Location data are provided in Data S1. Our final data set included a total of 487 occurrence records of all four diseases.

Information on 19 bioclimatic variables for each occurrence point was obtained from WORLDCLIM 2.0^70^ using the “extract” function in RASTER 3.0-7^71^. Mean values for each bioclimatic variable for each taxon are provided in Data S1.

### Climatic niche evolution

Method 1: We used the average bioclimatic conditions in a principal components analysis (PCA) based on their correlation matrix to illustrate climatic niche occupation of different diseases. The axes to be retained for further analyses were determined using the broken-stick method as implemented in VEGAN 2.5-6^72^. We assumed that the measured variant/strain means are a reasonable approximation of the realized climatic niche of the variant/strain^26,73^. However, given the fact that only a single geo-coordinate exists for most *Bd* isolates (except for *Bsal* and a few isolates of ranaviruses and Perkinsea), we caution that, there is a possibility of sampling artifacts or localized acclimatization can influence the calculation of mean bioclimatic variables from this data set. Therefore, we examined if calculated PC scores of each isolate/strain of each pathogen fall within the range of observed variation for the grouping considered. We used geographical regions as the common grouping variable for all four pathogens. Because most data points fall within the range of observed variation, we regard them as valid and used them in further analyses (Extended Data Figures 9-12). Outliers were not removed from the analyses, as they do not affect the interpretation of our results.

We assessed the extent to which traits (climatic niche evolution along each PC axis) had accumulated over time in each biogeographic region using disparity-through-time (DTT) plots^74^, with expected disparities calculated based on 1000 resamplings by using the ‘dtt’ function in GEIGER 2.0.6.2^75^ and with phenograms (projections of the phylogenetic tree in a space defined by phenotype/PC axis and time) constructed by using the ‘phenogram’ function in PHYTOOLS.

Although we wanted to test if climate is non-randomly associated with isolates of each lineage and their virulence, relevant data are available only for *Bd*. Therefore, we conducted this analysis only for *Bd* by following the classification of lineages and virulence of^5^. We traced the six main lineages of *Bd* (GPL, CAPE, ASIA-1, ASIA-2, CH and HYBRID) on the PCA plot as well as on the traitgrams (Fig.4). We also traced virulence and non-virulence on the PCA plot and the traitgram to see if phylogenetic signals associated with climate are visible in relation to virulence (Extended Data Figure 7).

Method 2: We used the ‘package nichevol 0.1.19^76^ in R 3.6.1 (<www.r-project.org>), an alternative method to check for evidence of climatic niche evolution, on the 19 bioclimatic variables for these diseases. This method enables critical steps for assessing ecological niche evolution over phylogenies, with uncertainty incorporated explicitly in reconstructions. It relies on ancestral reconstruction of ecological niches using a bin-based approach that incorporates uncertainty in estimations. Compared to other existing methods, this approach reduces the risk of overestimation of amounts and rates of ecological niche evolution^76^. However, since the current data set does not yield sufficient variance for occurrence records of most of the variants/isolates, we categorized all occurrence points into their strain or a clade. We categorized chytridiomycosis into its specific strains; Bd-GPL, CAPE, ASIA-1, ASIA-2 and Bsal (CH and HYBRID were omitted as they contain very few occurrence records). Ranavirus occurrences were categorized as ATV-like, CMTV-like and FV3-like, and Perkinsea occurrences were categorized into clades A, B, C and Pathogenic Perkinsea Clade (PPC). The phylogenetic trees were also pruned and renamed so that only the considered strains/clades were retained.

As spatial distribution polygons are not available for above strains or clades, we created spatial distribution polygons as ‘.shp’ files. We used all occurrence records of a variant/clade and predicted the potential geographical distribution of them using MaxEnt^32^; see the “ecological niche modeling” section below for more methodological details. This allowed us to define the fundamental niches of these strains/clades and to detect areas that present suitable conditions/accessible areas (termed **M**)^77^, which in turn are the areas across which niche models should be calibrated^78^.

We developed tables of character values for all strains/clades considered for each environmental dimension (19 bioclimatic variables), summarizing ranges of variable values occupied by each strain/clade and manifested across each species’ **M**. Variable values in M then were categorized into multiple classes (bins) and, by using values in occurrences, the presence or absence of the species in each bin was tested. Values of characters are 0 = absent, 1 = present, and ? = unknown (uncertain). Ancestral reconstructions of ecological niches (represented by 19 bioclimatic variables) were performed using both the MCC tree and 1000 posterior trees. Reconstructions were performed using maximum parsimony and maximum likelihood methods (results were more or less similar, but variation was higher in MP). Next, the variability of each bioclim variable was plotted on separate MCC trees (Extended Data Figures 4-6). Overall, this method allowed us to combine both NC (through MaxEnt based niche models) and CNE (through ancestral reconstructions) to understand their combined effects.

### Spatial point pattern analysis and ecological niche modeling

To assess the potential distribution of these four amphibian diseases, we mapped their climatic niches based on environmental layers and their global occurrences obtained above. The model was built using two methods (Extended Data Figure 1): the maximum entropy algorithm MaxEnt^32^ in the R packages ‘dismo’^71^, and ‘ENMeval’^79^ and spatial point process analysis (PPM)^31^ in the R package ‘spatstat’^80^. In the main text we only provide the results obtained from MaxEnt (Fig. 1) as results from both methods were very similar. Results obtained using PPMs are provided in supplementary material.

MaxEnt estimates the probability of species occurrence by finding the distribution of maximum entropy, which is subject to constraints defined by the environmental variables being analyzed. Nineteen bioclimatic layers were extracted from the WorldClim 2.1 database (http://www.worldclim.org) for present-day conditions (∼ 1970–2000) at a spatial resolution of 30 arc-seconds (∼ 1 km). Predictor collinearity was eliminated by calculating Pearsons’s correlation coefficients for all pairs of bioclimatic variables, excluding the variables from a correlated pair (|r| > 0.85. The MaxEnt model was optimized using the ENMeval package^79^ and the records were partitioned for background testing and training data to check for spatial autocorrelation and over-fitting. Model performance was measured using the Area Under the Curve (AUC) and the results were overlaid on raster maps.

Point process models (PPMs) are used to analyze species presence-only data in a regression framework^31,81^. Presence-only data typically arise as point events – a set of point locations where a species has been observed. In the statistical literature, a set of point events (in which the location and number of points is random) is known as a point process. However, PPMs are closely connected to methods already in widespread use in ecology such as MAXENT^82-84^, some implementations of logistic regression^31,85^ and estimation of resource selection functions^82,86^. Here we used both MaxEnt and PPMs because PPMs confer particular benefits for interpretation and implementation.

PPMs are a natural choice of analysis method for presence-only SDM, especially when the data arise as point events. Initially, we converted our raster covariates (19 bioclimatic variables at a spatial resolution of 2.5 arcmin) into image objects using as.im.RasterLayer function from the ‘maptools’ library. Next, our analysis window was set up and our point pattern object was created to the extent of the world map. After accounting for edge effect, we explored for evidence of spatial dependence using Ripley’s K and the more intuitive L function along with envelope tests. According to the Kest, Lest and envelope tests, our data deviated from complete spatial randomness. Therefore, an inhomogeneous point processes model was developed. We used the quadscheme function from spatstat to create quadrature points. The resolution of these points and the best-performing value for quadrate scheme was determined by setting the number of grid points in the horizontal and vertical directions and by checking for lowest AIC values, respectively. We experimented many combinations of predictor variables and tested their suitability through comparison of the model AIC statistics. As our initial model results exhibited spatial clustering, we used a Matern process to account for this. The model coefficients were checked by calling coef (maternMod) to see the relationship between our predictor variables and the predicted intensity of points. The AUC value, which is the probability that a randomly selected data point has higher predicted intensity than does a randomly selected spatial location, was calculated. The roc curve and auc statistic for our models were high, suggesting a well-specified inhomogeneous point process model. Finally, a mapped output of our results was generated using the predict function (Extended Data Figure 13).

For future projections, we used future climate change scenarios for the years 2021-2040 based on the CMIP6 downscaled future climate projections available at https://www.worldclim.org/data/cmip6/cmip6climate.html. We used nine global climate models (GCMs)—BCC-CSM2-MR, CNRM-CM6-1, CNRM-ESM2-1, CanESM5, GFDL-ESM4, IPSL-CM6A-LR, MIROC-ES2L, MIROC6 and MRI-ESM2-0—and four Shared Socio-economic Pathways (SSPs)—126, 245, 370 and 585—to make future projections using MaxEnt. Mean model values obtained were overlaid on raster maps to assess the future potential distribution of the four amphibian diseases. Finally, we measured the differences among present and future rasters for each disease using raster calculations in package ‘raster’ ^87^ in R v. 3.6.1^88^.

## Data and materials availability

The authors declare that data supporting the findings of this study are available within the file Data S1. All codes used in support of this publication and the additional data are publicly available in specific R packages and public data repositories as indicated in the text.

## Acknowledgments

We thank Tingru Mao for assistance with the data analysis; Guangxi University Laboratory Startup Funding (MM), Ziff Environmental Postdoctoral Fellowship, Harvard University Center for the Environment (MM), National Research Council Sri Lanka 11-124 (MM) and China Student Council Fellowship for graduate studies (GE, JH1) for providing financial assistance and the Wildlife Heritage Trust of Sri Lanka (RP).

## Author contributions

GE, JH1, SD and MM conceptualized the study. GE, JH1, SD, MRP, KAM, MM designed methodology. RP, JH2, MM acquisitioned funding. All authors contributed in writing the original draft, review and editing. MM supervised the study.

## Competing interests

Authors declare that they have no competing interests.

## Notes

### Competing Interest Statement

The authors have declared no competing interest.

