## Supplementary Information for "Climatic niche evolution of infectious diseases driving amphibian declines"

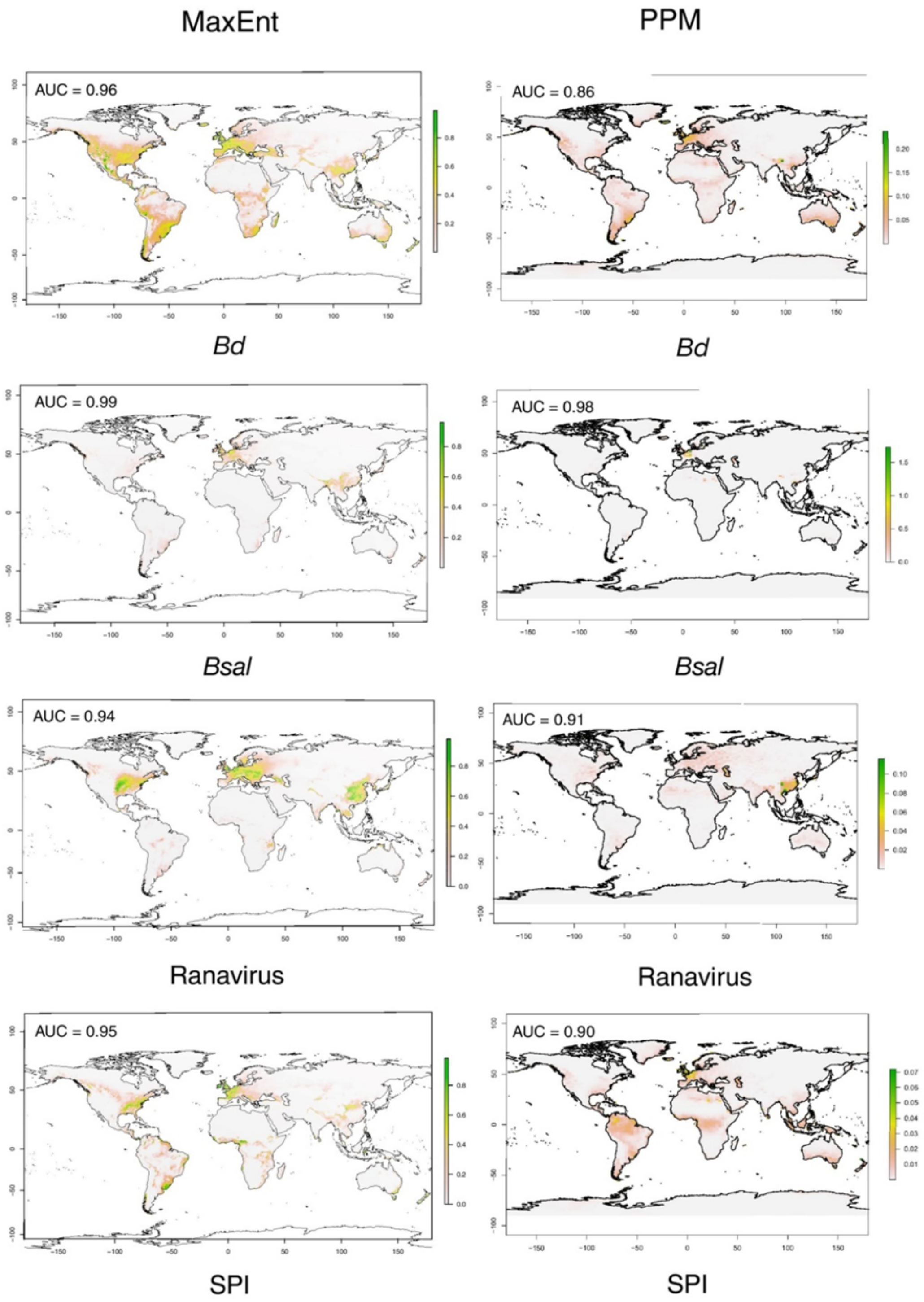


**Extended Data Figure 1. Comparison of results from MaxEnt species distribution modeling to those from spatial point process models (PPM).** Both analyses yielded more or less similar distribution ranges for all four diseases, although probability of occurrence is lower in the PPM analyses. AUC values are indicated on maps.

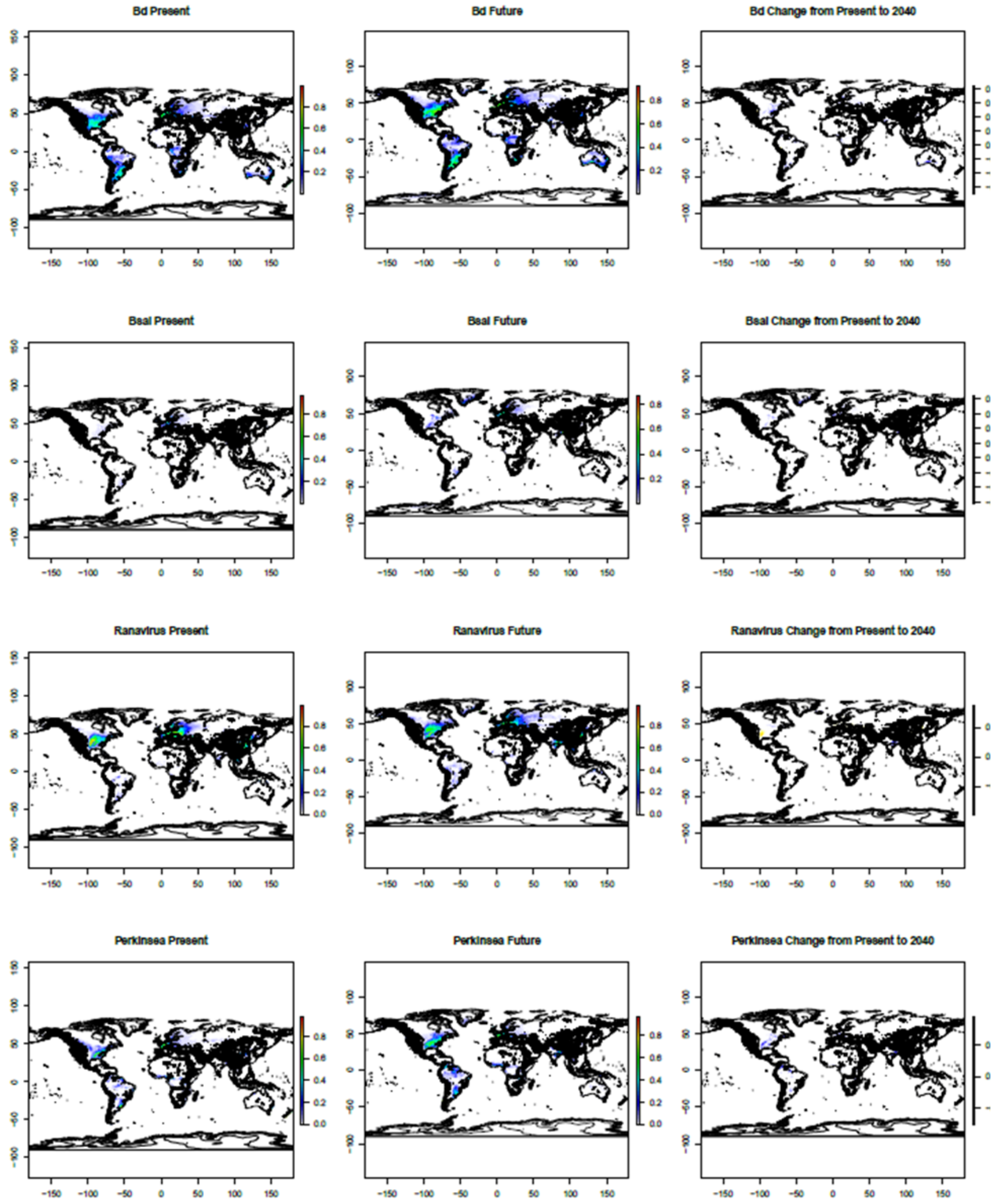


**Extended Data Figure 2. 1000-meter-elevation contour lines traced on potential distribution range expansion of *Bd*, *Bsal*, ranaviruses and Perkinsea two decades from now based on CMIP6 downscaled future climate projections.** *Bd* and *Bsal* exhibit potential expansion into higher elevations but ranaviruses and Perkinsea are more likely to expand in areas below 1000 m. See enlarged PDF for clarity.

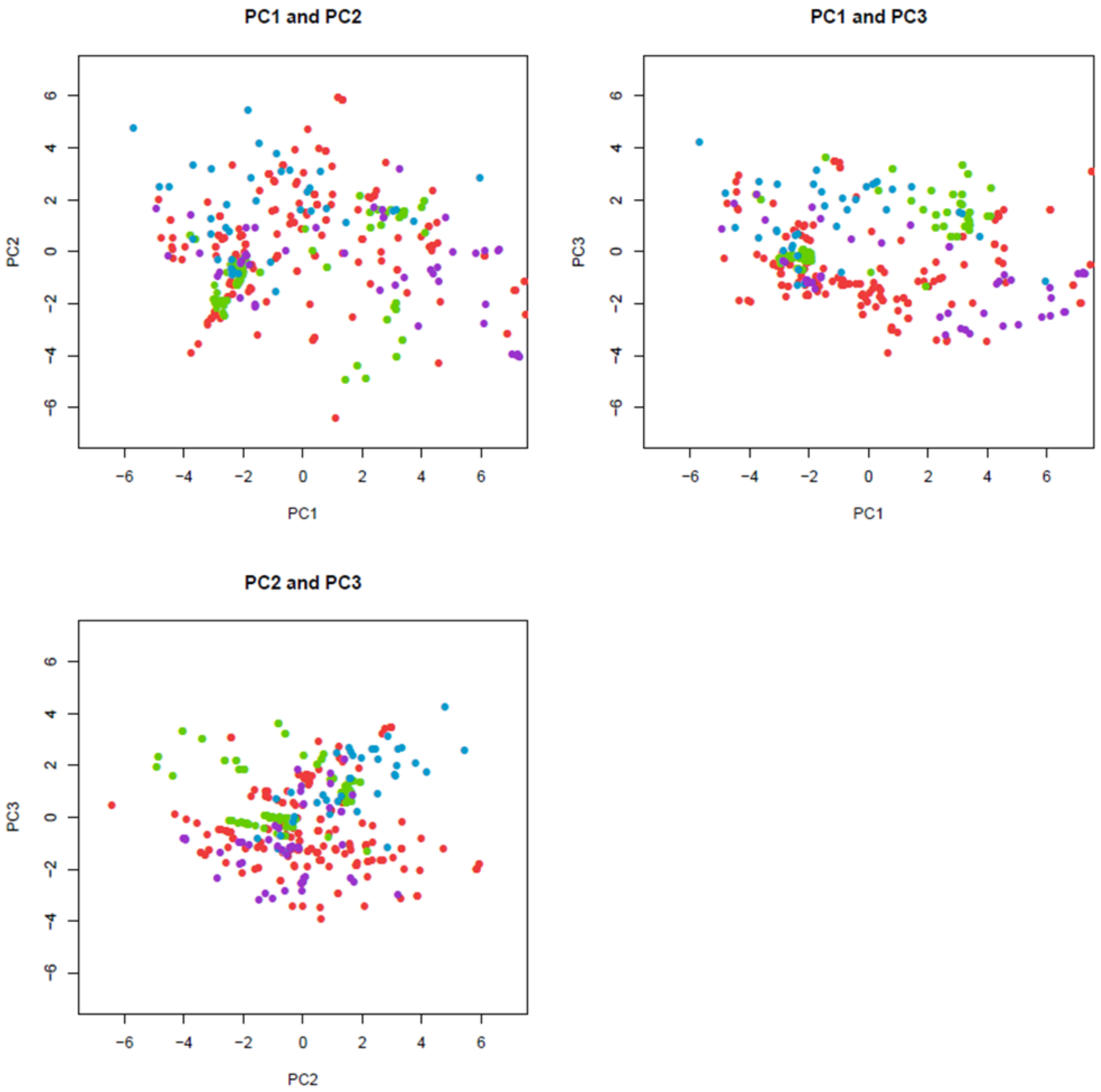


**Extended Data Figure 3. Climatic space defined by the first three principal component axes of climatic niches in amphibian diseases.** Different colors represent the four major amphibian diseases: *Bd* (red), *Bsal* (green), ranaviruses (purple) and Perkinsea (blue). Each point represents the average climatic conditions for each taxon. Loadings are provided in Supplementary Table 2. The first three PC axes explain over 75% of the variance in the data. Seasonality and precipitation variables segregate the climatic niche space of *Bsal* and Perkinsea from those of *Bd* and ranaviruses. Perkinsea and *Bsal* show positive PC3 values, which suggests dependence on precipitation seasonality, while *Bd* and ranavirus show negative values.

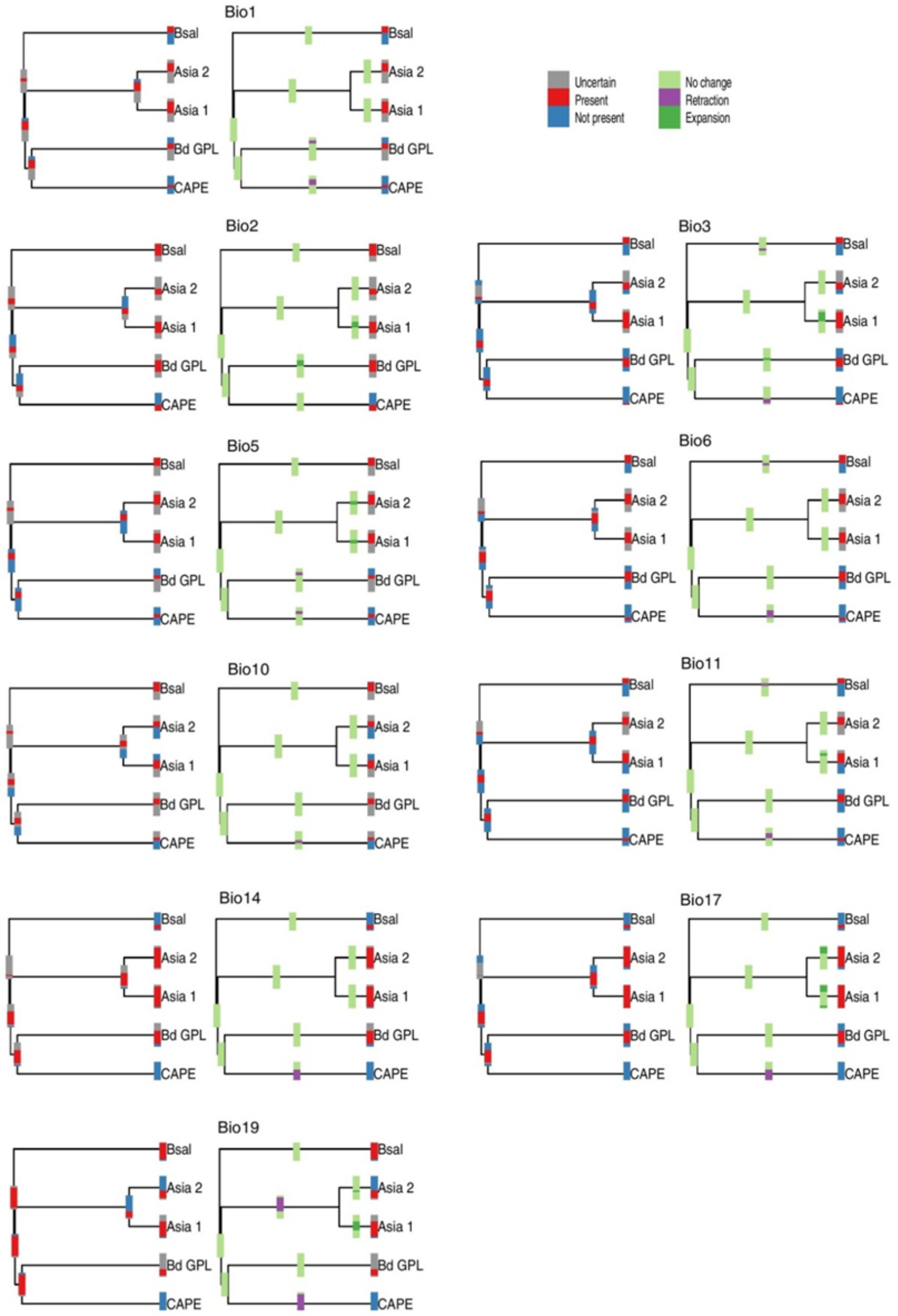


**Extended Data Figure 4. Maximum parsimony reconstructions of ecological niche evolution for strains of *Bd* and *Bsal* using the 10 environmental variables that dominate the first three principal component axes of their climatic niche space: bio1, bio2, bio3, bio5, bio6, bio10, bio11, bio14, bio17 and bio19.** Evidence of niche retraction by CAPE is seen in all variables. ASIA-1 shows niche expansion in bio2, bio3, bio5, bio11, bio17 and bio19, while *Bsal* shows niche retraction in bio3, bio6 and bio11.

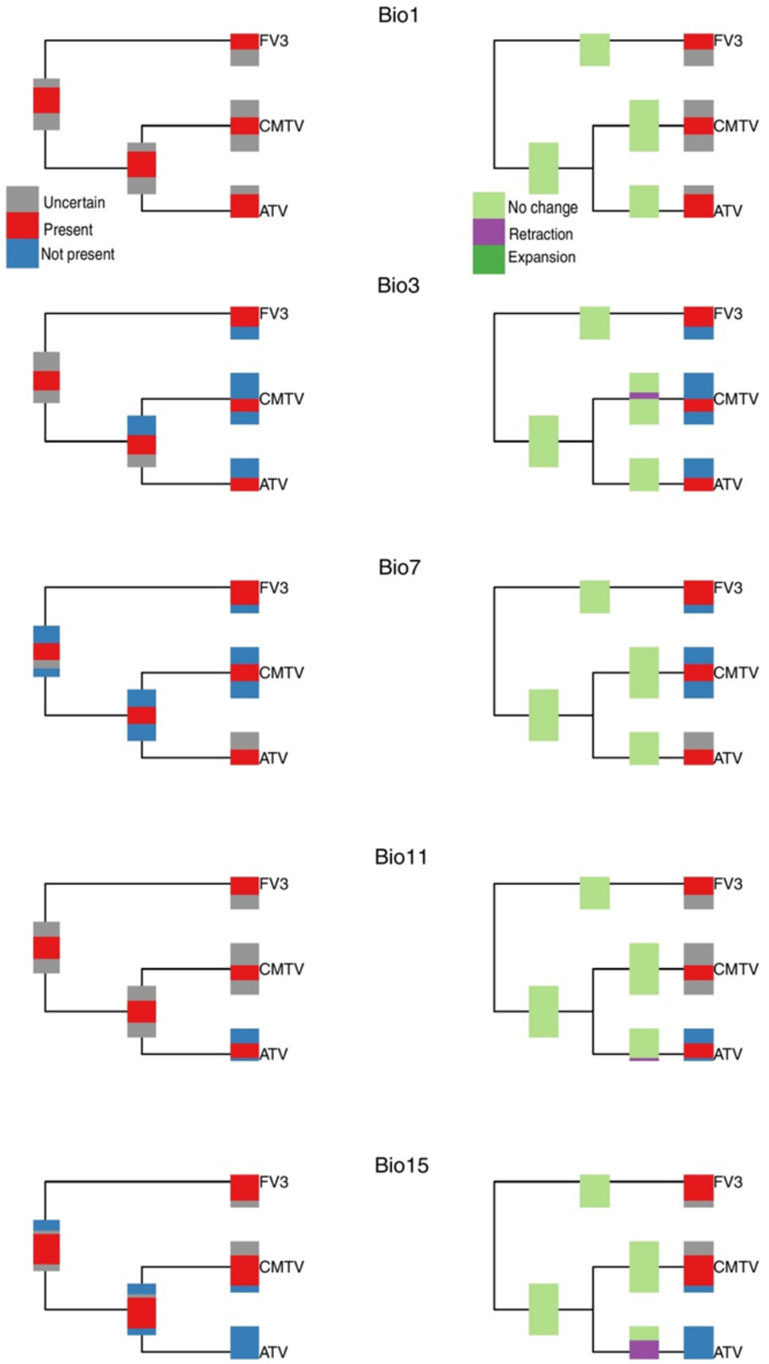


**Extended Data Figure 5. Maximum parsimony reconstructions of ecological niche evolution for major ranaviral strains using the five environmental variables that dominate the first three principal component axes of their climatic niche space: bio1, bio3, bio7, bio11 and bio15.** Evidence of niche retraction is shown for ATV-like ranaviruses in bio11 and bio15, and for CMTV-like ranaviruses in bio3. Niche expansion is evident for CMTV-like ranaviruses in bio6, bio14 and bio17 (not depicted).

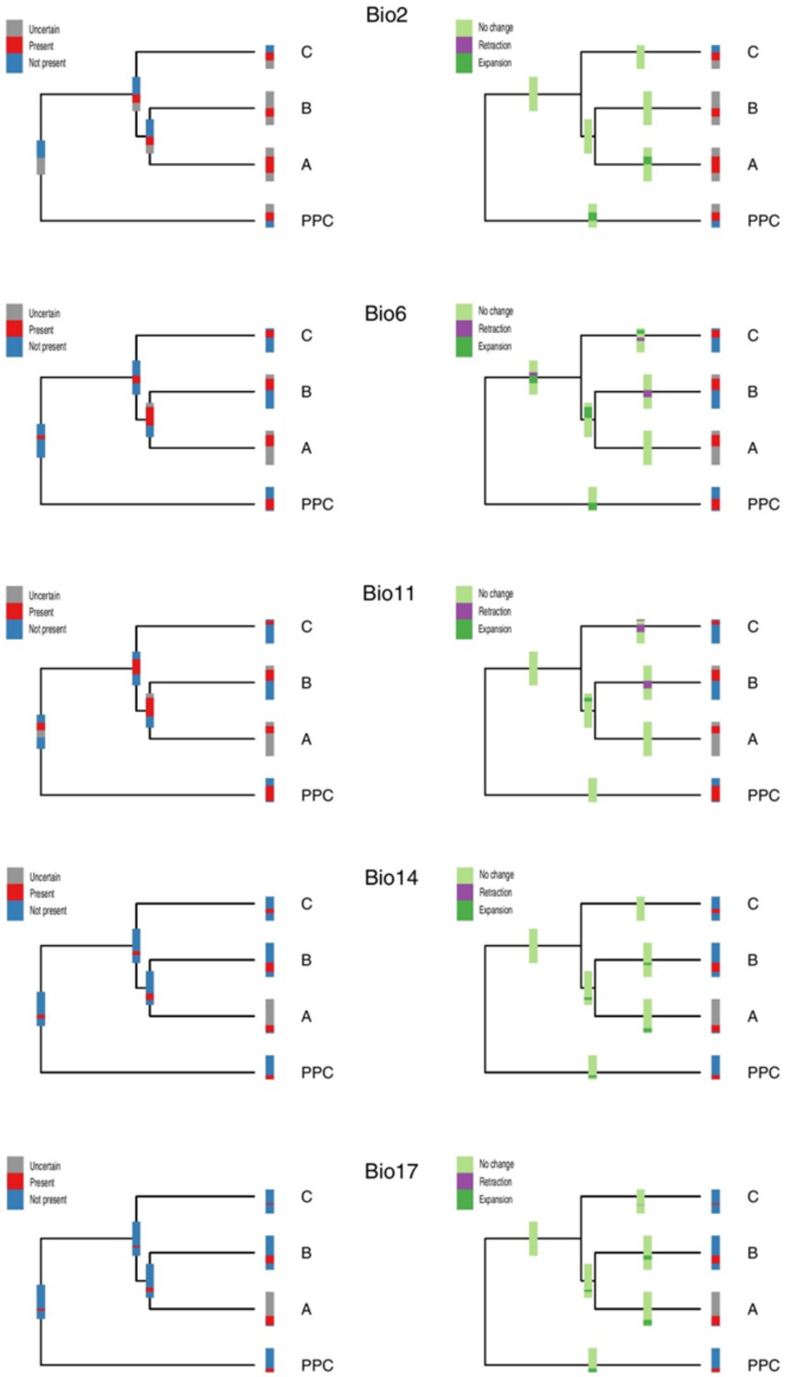


**Extended Data Figure 6. Maximum parsimony reconstructions of ecological niche evolution for major SPI clades using the five environmental variables that dominate the first three principal component axes of their climatic niche space: bio2, bio6, bio11, bio14 and bio17.** Evidence of niche expansion for the Pathogenic Perkinsea Clade (PPC) is shown along bio2, bio6, bio14 and bio17, and for Clade A along bio2, bio14 and bio17. Niche retraction for clades B and C are evident along bio6 and bio11.

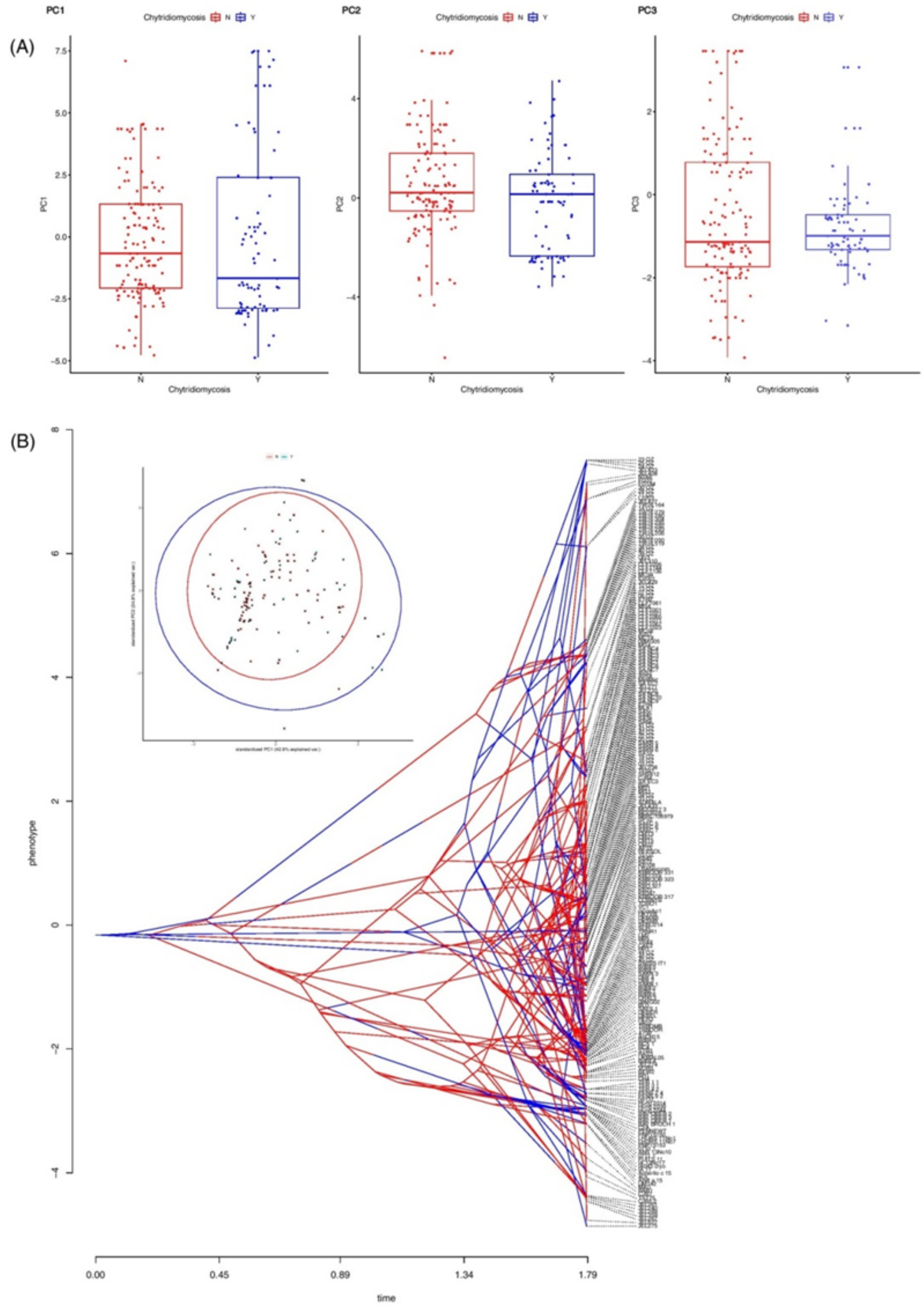


**Extended Data Figure 7. Climate correlates of virulence in *Bd* isolates.** (A) Boxplots showing PC1 deviation among virulent (red) and non-virulent (blue) isolates of *Bd*. A significant difference between PC1 scores for virulent and non-virulent *Bd* isolates is not observed. (B) Traitgram showing climatic niche evolution in virulent vs. non-virulent strains of *Bd*. A distinct phylogenetic signal is not observed between the two categories. Inset shows climatic niche occupation of virulent vs. non-virulent isolates of *Bd*. Virulent isolates have a broader climatic niche occupation than non-virulent ones.

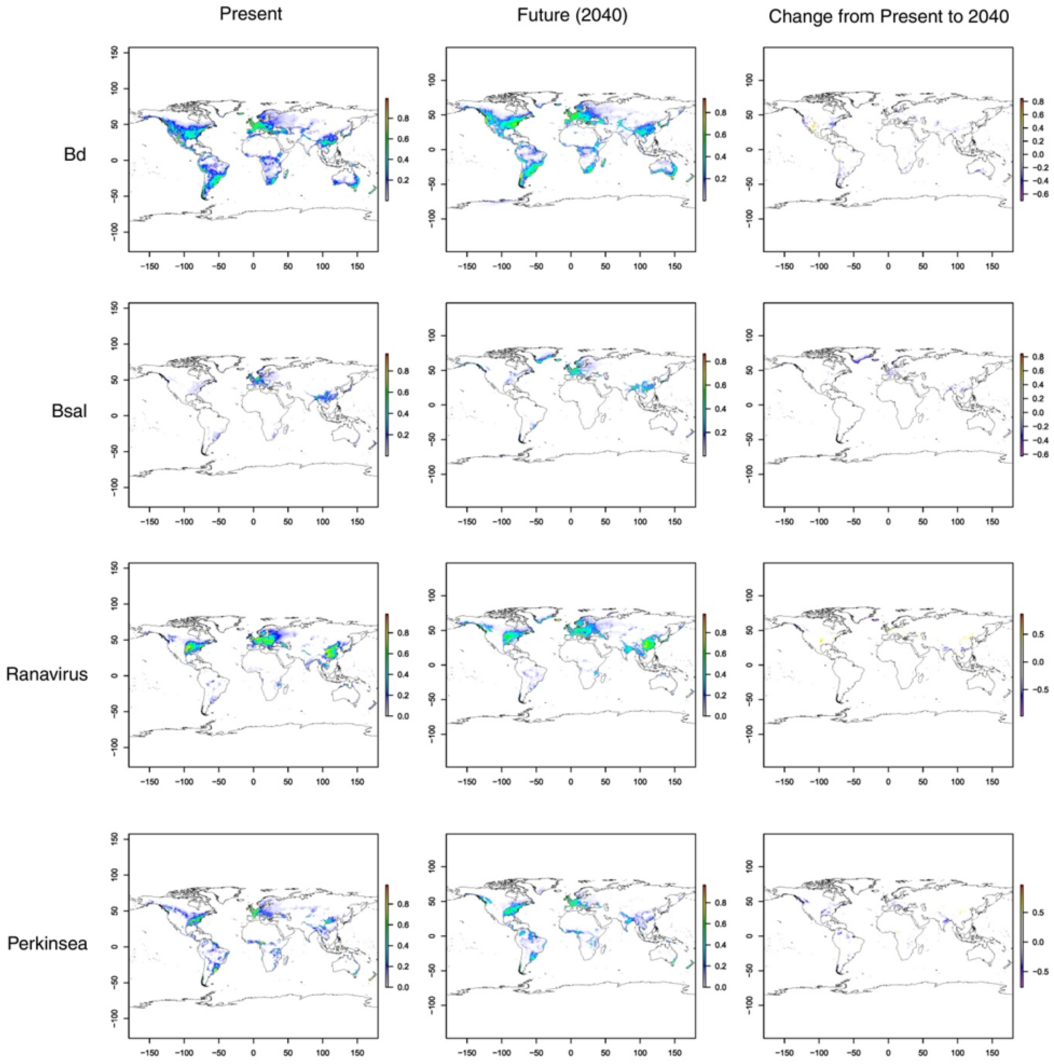


**Extended Data Figure 8. Present and predicted future (2040) distributions, and resulting range expansion, of *Bd*, *Bsal*, ranaviruses and Perkinsea based on CMIP6 downscaled future climate projections.** Potential distributions of *Bd*, *Bsal*, ranavirus and Perkinsea under current climatic conditions (left column) are based on SDM. Potential distribution of the diseases under future climatic scenarios (2021–2040 under CMIP6 downscaled future climate projections; middle column). Mean range expansion from present to 2040 (right column). Note the future reduction of suitable areas for *Bd* and *Bsal* and the possible expansion of *Bd* into North America, South America, parts of South Africa and Sunda islands. An expansion of suitable habitats for ranaviruses is evident throughout the tropics and subtropics in regions such as North America and Europe. Perkinsea too will expand into Asia, to Africa, Panama isthmus and some parts in Indonesia in another 20–30 years. Warmer colours indicate higher or increased suitability; cooler colours indicate lower or decreased suitability (see enlarged PDF for clarity).

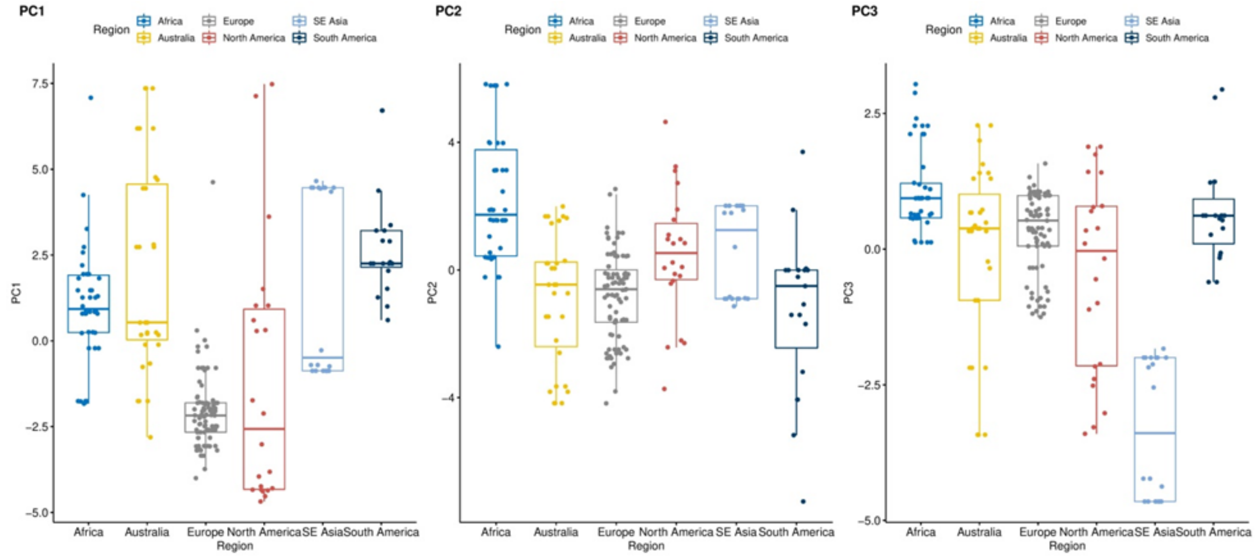


**Extended Data Figure 9. Boxplots showing the data distribution of calculated PC scores for each *Bd* isolate categorized based on geographical region.** Each plot depicts mean, quartiles and standard deviation. Most data points fall within the range of observed variation for the grouping considered.

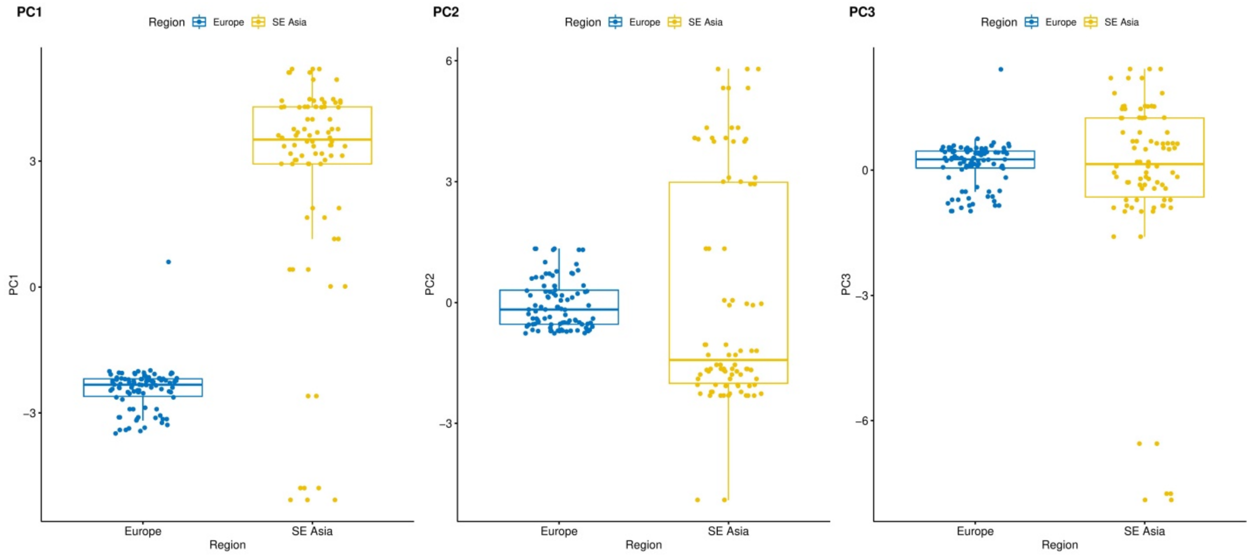


**Extended Data Figure 10**. **Boxplots showing the data distribution of calculated PC scores for each *Bsal* isolate categorized based on geographical region.** Each plot depicts mean, quartiles and standard deviation. Most data points fall within the range of observed variation for the grouping considered.

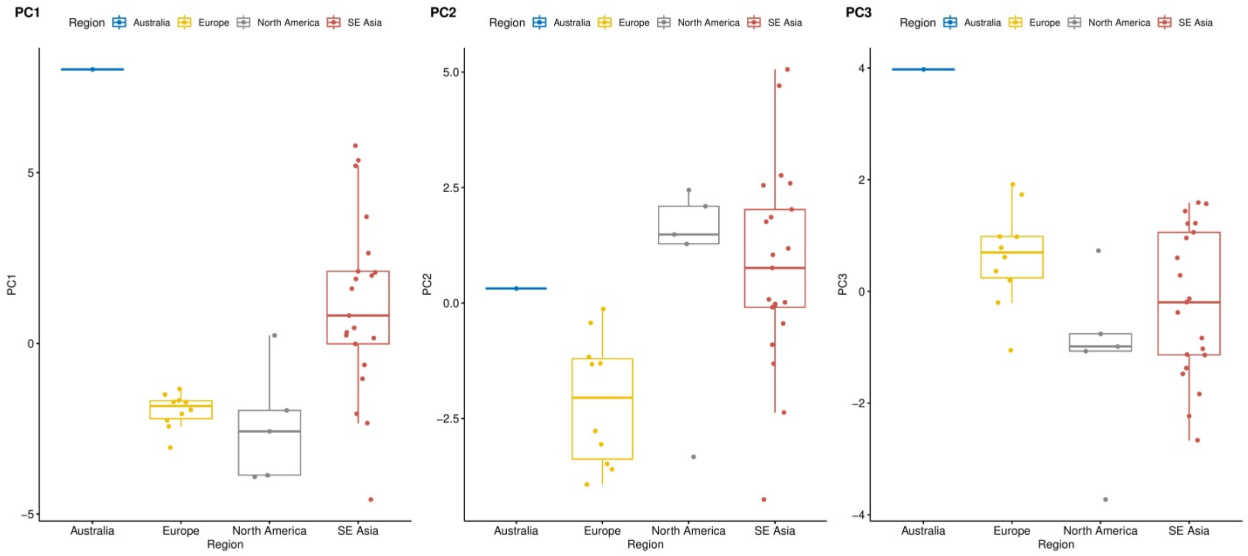


**Extended Data Figure 11. Boxplots showing the data distribution of calculated PC scores for each ranaviral strain categorized based on geographical region.** Each plot depicts mean, quartiles and standard deviation. Most data points fall within the range of observed variation for the grouping considered.

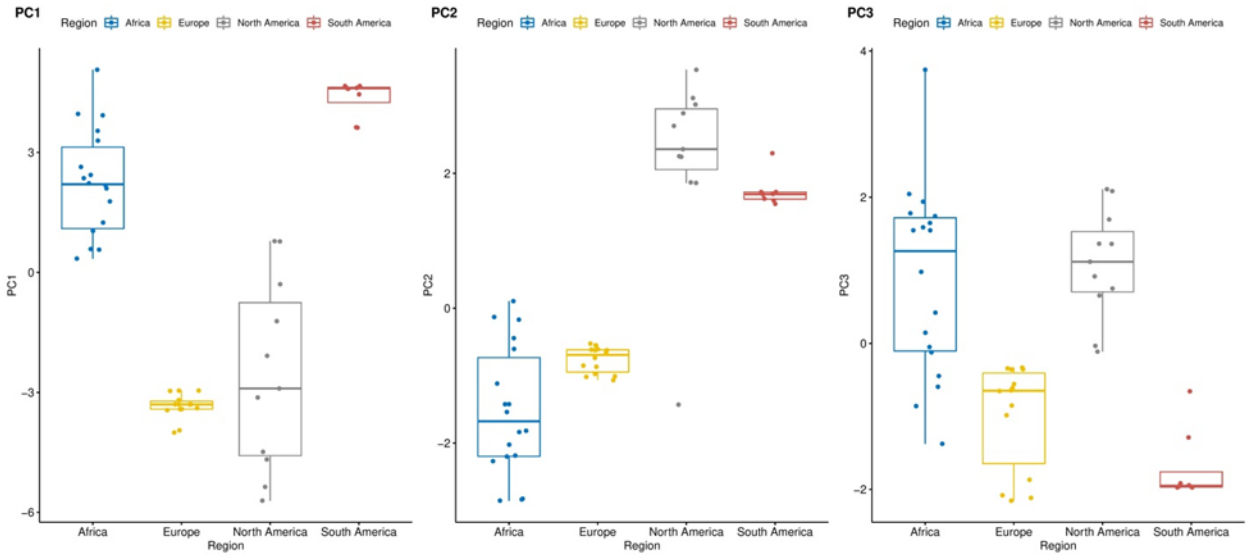


**Extended Data Figure 12. Boxplots showing the data distribution of calculated PC scores for each Perkinsea isolate categorized based on geographical region.** Each plot depicts mean, quartiles and standard deviation. Most data points fall within the range of observed variation for the grouping considered.

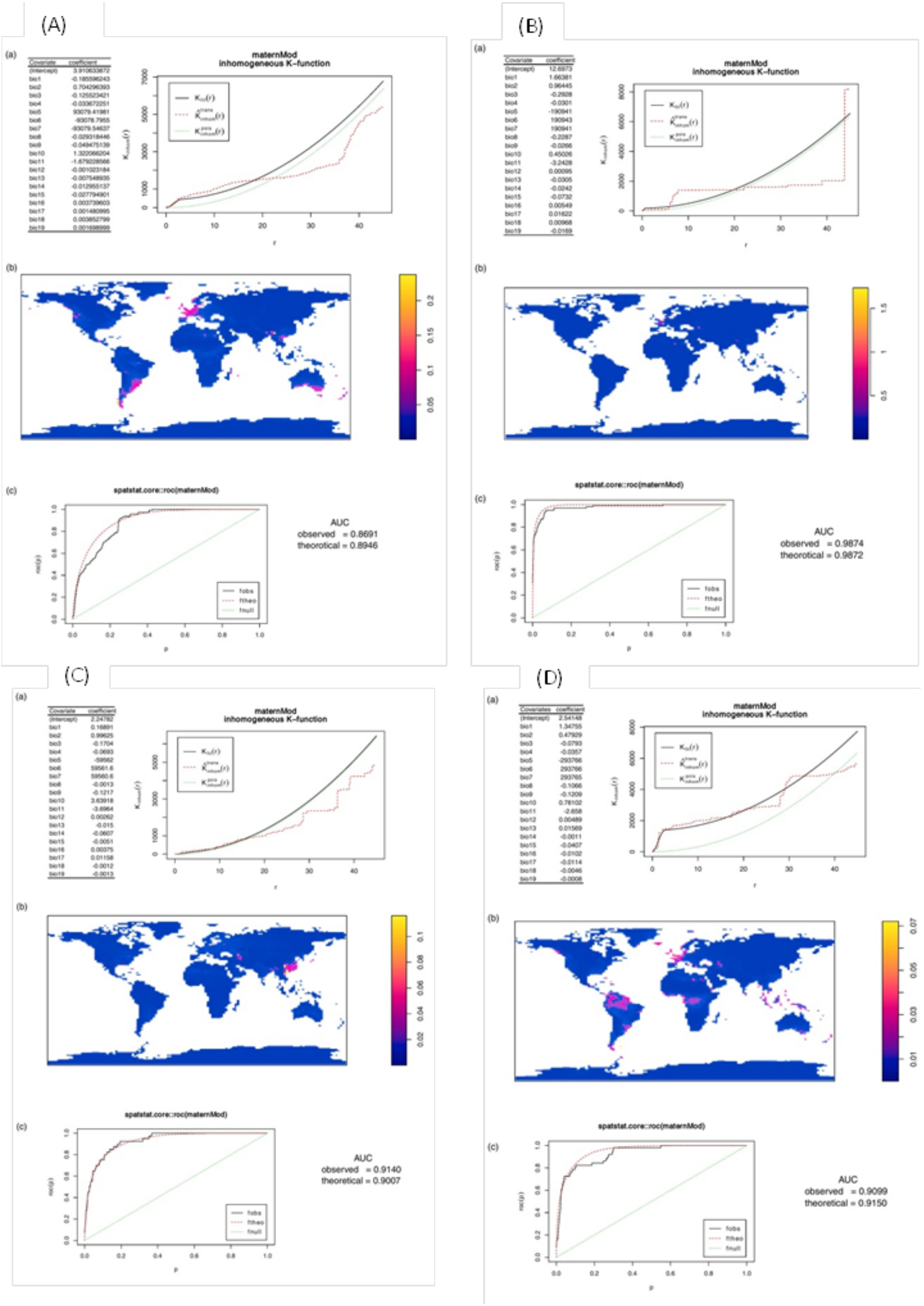
(A)
(a)
Covariate coefficient
(Intercept) 3.910633872
bio1 -0.185596243
bio2 0.704296393
bio3 -0.125523421
bio4 -0.033672251
bio5 93079.41981
bio6 -93078.7955
bio7 -93079.54637
bio8 -0.029318446
bio9 -0.049475139
bio10 1.322066204
bio11 -1.679228566
bio12 -0.001023184
bio13 -0.007548935
bio14 -0.012955137
bio15 -0.027794901
bio16 0.003739603
bio17 0.001480995
bio18 0.003852799
bio19 0.001698999
maternMod
inhomogeneous K-function
(b)
(c)
spatstat.core::roc(maternMod)
AUC
observed = 0.8691
theorotical = 0.8946
(B)
(a)
Covariate coefficient
(Intercept) 12.6973
bio1 1.66381
bio2 0.96445
bio3 -0.2928
bio4 -0.0301
bio5 -190941
bio6 190943
bio7 190941
bio8 -0.2287
bio9 -0.0266
bio10 0.45026
bio11 -3.2428
bio12 0.00095
bio13 -0.0305
bio14 -0.0242
bio15 -0.0732
bio16 0.00549
bio17 0.01622
bio18 0.00968
bio19 -0.0169
maternMod
inhomogeneous K-function
(b)
(c)
spatstat.core::roc(maternMod)
AUC
observed = 0.9874
theorotical = 0.9872
(C)
(a)
Covariate coefficient
(Intercept) 2.24782
bio1 0.16891
bio2 0.99625
bio3 -0.1704
bio4 -0.0693
bio5 -59562
bio6 59561.6
bio7 59560.6
bio8 -0.0013
bio9 -0.1217
bio10 3.63918
bio11 -3.6964
bio12 0.00262
bio13 -0.015
bio14 -0.0607
bio15 -0.0051
bio16 0.00375
bio17 0.01158
bio18 -0.0012
bio19 -0.0013
maternMod
inhomogeneous K-function
(b)
(c)
spatstat.core::roc(maternMod)
AUC
observed = 0.9140
theoretical = 0.9007
(D)
(a)
Covariates coefficient
(Intercept) 2.54148
bio1 1.34755
bio2 0.47929
bio3 -0.0793
bio4 -0.0357
bio5 -293766
bio6 293766
bio7 293765
bio8 -0.1066
bio9 -0.1209
bio10 0.78102
bio11 -2.658
bio12 0.00489
bio13 0.01569
bio14 -0.0011
bio15 -0.0407
bio16 -0.0102
bio17 -0.0114
bio18 -0.0046
bio19 -0.0008
maternMod
inhomogeneous K-function
(b)
(c)
spatstat.core::roc(maternMod)
AUC
observed = 0.9099
theoretical = 0.9150

**Extended Data Figure 13. Point process models fitted to point pattern datasets of the four amphibian diseases: (A) *Bd*; (B) *Bsal*; (C) ranaviruses; and (D) Perkinsea.** In (A), each figure indicates model coefficients obtained from the full point process models. The covariates used in the final model were selected based on the relative contribution of each covariate. The figure on the right shows the behavior of the final inhomogeneous cluster point process model developed. The inhomogeneous K function (solid black line), the K function under trans edge correction method (red dotted line) and the theoretical inhomogeneous K function under the null hypothesis that the points are completely randomly distributed (green dashed line) are indicated. Where the observed K exceeds the theoretical K, we see more clustering than expected, and where the observed K is less than the theoretical K we see more dispersion than expected. In (B), maps depict outputs of the developed inhomogeneous cluster point process models. In (C), AUC values for each model are provided. Here, AUC is the probability that a randomly selected data point has higher predicted intensity than does a randomly selected spatial location.

### Supplementary Tables

### Supplementary Table 1. Summary of the prominent characteristics of the diseases

| Characteristics | Ranaviruses | *Batrachochytrium dendrobatidis (Bd)* | *Batrachochytrium salamandrivorans (Bsal)* | *Perkinsea/Severe Perkinsea infection (SPI)* | Reference |
|---|---|---|---|---|---|
| Variants/Strains | *Ambystoma tigrinum virus* (ATV), *Common midwife toad virus* (CMTV), *European North Atlantic ranavirus* (LfRV), *Epizootic haematopoietic necrosis virus* (EHNV), *Frog virus 3* (FV3), *Santee-Cooper ranavirus and Singapore grouper iridovirus* (SGIV)<br>Ranavirus isolates are considered members of the same viral species if they share > 95% amino acid identity.<br>*Ranavirus maximus, cod iridovirus* and *short-finned eel virus* are potential new species that remain unclassified. | Lineages<br>*Bd*CH,<br>*Bd*ASIA-1,<br>*Bd*ASIA-2/BRAZIL, *Bd*ASIA-3,<br>*Bd*GLP,<br>*Bd*CAPE,<br>Hybrids | Currently exists as a single lineage | Different NAG01 Perkinsea lineages in different continents (e.g., Pathogenic Perkinsea Clade-PPC in North America). | [3,5,18,66,89] |
| Host range | Ectothermic vertebrates (fishes, amphibians and reptiles). | All orders in Amphibia and few non-amphibans, e.g., crayfish | Urodela in Amphibia | Frogs in Anura | [3,8,86,90] |

| | | | | | |
|---|---|---|---|---|---|
| No. of host species / classes | At least 175 species across 52 families of ectothermic vertebrates (by 2015); at least 105 species of amphibians in 18 families belong to 46 genera (by 2015). | 1375 amphibian species has been detected by *Bd* | Over 60 urodele species | 20 genera/phylogroups from five countries across three continents | 3,7,8,10,16,18,40,68,91-100 |
| Infect wild populations/ cultured species/ Introduced species | Wild (native, endemic), cultured and introduced | Wild (native, endemic), cultured and introduced | Wild (native, endemic), cultured and introduced | Infect wild populations/ cultured species/ introduced species | 5,7,18 |
| Mode of transmission | Close contact, feeding infected individuals, via water | Direct physical contact or indirectly via spores in media | Direct physical contact or indirectly via spores in media | Direct physical contact or indirectly via spores in media | 41,101,102 |
| Time of evolution/ emerging as pathogen | Unknown | Highly pathogenic *Bd*GLP lineage evolved between 120 and 50 years ago | Unknown | Unknown | 5 |
| Time of first discovery and location | FV3 was discovered in the 1960s from USA<br>Host: northern leopard frogs (*Lithobates pipiens*) | Bd was described in 1999 in USA due to the mortality of frogs found in the 1900s in Central America and Australia | *Bsal* was described in 2013 in The Netherlands | 1999 in North America | 13-15,35 |
| How they evolved from ancestry | Unknown | Trade across continents | Trade across continents | Unknown | 5,7 |
| Currently inhabiting regions | All continents except Antarctica | All continents except Antarctica | Europe and Asia | North and South America, Europe, and Africa | 3,7,8,16,18,90,103 |
| Climatic/seasonal/ environmental preference | In temperate areas, ranaviruses generally cause outbreaks with amphibian mass mortality in the summer months. | Under experimental conditions, survive between17 and 25℃, field data often show seasonal patterns in prevalence, corresponding to more favorable climatic conditions; e.g., cooler seasons in the tropics, where high temperatures may limit the pathogen during other parts of the year and warmer seasons in temperate areas or at high elevations, | Under experimental conditions, survive within relatively cooler thermal limits (10–15℃), optimal temperatures | Outbreaks occur from summer to early autumn in boreal and temperate regions, and from late winter to early spring (and occasionally during the summer) in | 14,18,35,39,104-106 |

| | | | | | |
|---|---|---|---|---|---|
| | | where low temperatures limit the pathogen for part of the year, and warmer seasons in temperate areas or at high elevations, where low temperatures limit the pathogen for part of the year. | being lower than *Bd* | regions with a subtropical climate | |
| Pathogenicity/ Intensity/Mortality rates | High intensity and mortality in most host species. Mortality reaches100% where they occur. | Highly virulent/high intensity/rapid mortality | Highly virulent/high intensity/rapid mortality | Highly virulent/high intensity/mortality rates as high as 95% | [7,16,107] |
| Any extinction of species caused by the disease | Not yet | Yes | Not yet | Not yet | [6] |
| Whether notifiable disease by World Organization for Animal Health (OIE ) | Infection by ranavirus species is listed as amphibian disease | A notifiable amphibian disease | A notifiable amphibian disease | Not yet | [12] |
| Factors favoring transmission | Generally summer disease. Rising global temperatures may favor the transmission. Global animal trade also facilitates the transmission. | Host species<br>Vectors<br>Climate<br>Habitat conditions<br>Relevant human activities | Host species<br>Vectors<br>Cooler climate<br>Habitat conditions<br>Relevant human activities | Host species<br>Water quality conditions<br>Relevant human activities | [14,18,47,108] |
| Control of the disease | Systematic surveillance, screening before trading of animals, regulations and monitoring in animal trade, vaccines are on trial | Biocontrol with predators, habitat bioaugmentation, enhance regulations in animal trade and long-term monitoring | Limiting amphibian trade, improving biosecurity measures, and increasing host resistance | Enhance disease monitoring in breeding season | [18,109,110] |

**Supplementary Table 2. Loadings of the principal components analysis of bioclimatic data associated with the occurrence of four amphibian diseases.** PC1, PC2 and PC3 explain 45.7%, 20.5% and 13.1% of the variance, respectively. PC1 is mostly associated with variation in temperature (bio1 and bio11) and PC2 with rainfall and temperature (bio2, bio5, bio15), whereas PC3 is associated with seasonal rainfall (bio4, bio18)**.**

| Bioclim variable | Description | PC1 | PC2 | PC3 |
|---|---|---|---|---|
| bio4 | Temperature Seasonality (standard deviation × 100) | -0.23 | 0.16 | 0.38 |
| bio18 | Precipitation of Warmest Quarter | 0.21 | -0.03 | 0.38 |
| bio7 | Temperature Annual Range (bio5-bio6) | -0.21 | 0.27 | 0.26 |
| bio8 | Mean Temperature of Wettest Quarter | 0.23 | 0.2 | 0.23 |
| bio13 | Precipitation of Wettest Month | 0.28 | -0.1 | 0.23 |
| bio16 | Precipitation of Wettest Quarter | 0.28 | -0.11 | 0.22 |
| bio12 | Annual Precipitation | 0.25 | -0.26 | 0.21 |
| bio10 | Mean Temperature of Warmest Quarter | 0.24 | 0.22 | 0.2 |
| bio14 | Precipitation of Driest Month | -0.05 | -0.4 | 0.18 |
| bio17 | Precipitation of Driest Quarter | -0.02 | -0.42 | 0.18 |
| bio5 | Max Temperature of Warmest Month | 0.21 | 0.28 | 0.13 |
| bio15 | Precipitation Seasonality (Coefficient of Variation) | 0.22 | 0.27 | 0.09 |
| bio1 | Annual Mean Temperature | 0.31 | 0.12 | 0.01 |
| bio19 | Precipitation of Coldest Quarter | 0.11 | -0.33 | -0.11 |
| bio11 | Mean Temperature of Coldest Quarter | 0.32 | 0.02 | -0.14 |
| bio6 | Min Temperature of Coldest Month | 0.31 | -0.05 | -0.14 |
| bio2 | Mean Diurnal Range (Mean of monthly (max temp - min temp)) | -0.01 | 0.34 | -0.21 |
| bio9 | Mean Temperature of Driest Quarter | 0.27 | -0.02 | -0.24 |
| bio3 | Isothermality (bio2/bio7) (× 100) | 0.22 | 0.01 | -0.39 |
| Total | | 45.7 | 20.5 | 13.1 |
| Cumulative | | 45.7 | 66.2 | 79.3 |

**Supplementary Table 3. Contributions of climatic variables in each PC axis for each disease.** The climatic niches of the four diseases differ along the PC2 and PC3 axes. Temperature variables are represented on PC1 for all diseases. *Bsal* and Perkinsea seem to be dependent on rainfall during the driest periods, whereas *Bd* and ranaviruses seem to be less dependent on rainfall on PC2 but require isothermal conditions along PC3.

| PC axis | *Bd* | *Bsal* | *Ranavirus* | Perkinsea |
|---|---|---|---|---|
| **PC1** | Annual mean temperature, Mean temperature of coldest quarter, Minimum temperature of coldest month | Annual mean temperature, Mean temperature of coldest quarter, Mean temperature of coldest month | Annual mean temperature, Mean temperature of coldest quarter | Mean temperature of coldest quarter, Minimum temperature of coldest month |
| **PC2** | Mean diurnal range | Precipitation of driest quarter, Precipitation of driest month, Precipitation of coldest quarter | Precipitation seasonality, Temperature annual range | Precipitation of driest month, Precipitation of driest quarter |
| **PC3** | Isothermality | Max temperature of warmest month, Mean temperature of warmest quarter | Isothermality | Mean diurnal range |

**Supplementary Table 4. Summary of the mitigating techniques for diseases, that can mostly be used *in vitro***

| Technique | Targeted disease | Targeted taxon | Effectiveness | Constraints | Reference |
|---|---|---|---|---|---|
| Using antifungal agents (Amphotericin B and voriconazole) | Chytridiomycosis *Bd* | Amphibians | Eliminated the *Bd* infection from all treated animals | Tested on captive breeding colonies. Effectiveness in natural occurring colonies may vary | [111] |
| Using antifungal agents (Sodium chloride) | Chytridiomycosis *Bd* | Amphibians | Significantly reduced the growth and motility of the chytrid fungus | Tested on captive breeding colonies. Effectiveness in natural occurring colonies may vary. There may be negative effects of salt exposure on both target and non-target organisms | [112] |
| Treating with protective bacteria (probiotics consortia) | Chytridiomycosis *Bd* | Amphibians | Bacterial consortia can offer stronger protection against *Bd* compared to single strains | Effect was not uniform across all *Bd* isolates | [113] |
| Treating with protective bacteria (probiotics consortia) | Chytridiomycosis *Bd* | Amphibians | Strong growth inhibition between 70 and 100% for seven Bd isolates | Two *Bd* isolates appeared consistently resistant to inhibition | [114] |
| Breeding more resistant individuals | Chytridiomycosis *Bd* | Amphibians | Increase disease resistance and mitigate the significant threat of chytridiomycosis | There might be undesired traits | [115] |
| DNA Vaccination | Ranavirosis | Salamanders | Significantly suppressed the virus replication. Conferred effective protection against ADRV infection | There might be practical implications to use among wild populations | [116] |
| DNA vaccine (encoding viral membrane protein) | Ranavirosis | Fish | Suppressed the virus replication and could induce protective immunity | There might be practical implications to use among wild populations | [117] |
